# Computational design of membrane fusion proteins

**DOI:** 10.64898/2026.05.04.722779

**Authors:** Masaharu Somiya, Sydney Funk, Dane Zambrano, Takumi Yanase, Noriyuki Hamaoka, Alex Kang, Banumathi Sankaran, Asim K. Bera, Neil P. King

## Abstract

The fusion of two distinct biological membranes is an evolutionarily conserved process essential to cellular organization and physiology. Membrane fusion is driven by the refolding of fusogenic proteins into low-energy postfusion states that overcome the energetic barrier to bilayer merger. Here we report a computational method for the design of synthetic fusogens inspired by the architecture of the soluble N-ethylmaleimide-sensitive factor attachment protein receptor (SNARE) complex. Using machine learning-guided protein design to extensively remodel backbone geometry and sequence, we generated heterodimeric SNARE-like assemblies that efficiently catalyze cell-cell membrane fusion. These minimal two-component fusogens exhibit substantially higher fusion activity than native multisubunit SNARE complexes. Structural and functional analyses identify the key determinants required for fusogenic activity and reveal a modularity that enables control of fusion through chemically induced heterodimerization. In addition to cell-cell fusion, the synthetic fusogens drive fusion between endoplasmic reticulum and mitochondrial membranes from human cells, demonstrating their potential as tools for programmable manipulation of intracellular membranes. Together, these results establish a general framework for the rational design of synthetic fusogens and expand the toolkit for engineering membrane dynamics in living systems.

## Introduction

Membrane fusion is a fundamental biophysical process in which two lipid bilayers merge to form a single continuous membrane. This process underlies a wide range of biological events, including fertilization (*1*), exocytosis (*2*), and viral entry (*3*). Although membrane fusion is thermodynamically favorable, it is kinetically constrained by a substantial energy barrier that must be overcome to progress from close membrane apposition through hemifusion stalk formation and, ultimately, fusion pore opening (*4*). In living systems, uncontrolled membrane fusion would be deleterious, as lipid bilayers are essential for cellular integrity and the compartmentalization required for homeostasis. Accordingly, membrane fusion is tightly regulated by specialized membrane fusion proteins, or fusogens, which catalyze bilayer merger by coupling membrane remodeling to energetically favorable protein conformational changes. Fusogens are found across all domains of life, including eukaryotes (*5–7*), archaea (*8*), enveloped viruses (*9*, *10*), and even non-enveloped viruses (*11*).

The biological importance of membrane fusion and its potential utility in biotechnological applications have inspired efforts to create synthetic membrane fusion machinery. For example, enveloped viruses rely on fusogens to efficiently merge viral and host membranes and deliver their genetic material, highlighting how synthetic fusogens could be useful for biologics delivery and gene therapy (*12*). However, due to the structural and mechanistic complexity of naturally occurring fusogens, most efforts to date have focused on simplified models of natural systems. Synthetic membrane fusion machineries constructed from complementary strands of DNA (*13–15*) or coiled-coil forming peptides (*16*, *17*) have provided valuable mechanistic insight into the physical principles underlying membrane fusion. Yet, because these systems have not been directly compared with efficient, naturally occurring fusogens, it is difficult to assess their potential biomedical utility. Furthermore, the limited set of viral fusogens explored in preclinical studies may be constrained by the risk of eliciting neutralizing anti-drug antibodies, which would preclude repeated administration (*18*). Thus there is a need to develop generalizable methods for designing synthetic protein-based fusogens with high activity.

Here we leveraged recent advances in deep learning-based methods for protein structure prediction and design to create streamlined heterodimeric SNARE-like fusogens with superior fusion activity compared to naturally occurring SNARE complexes. In addition to their potential biomedical utility, our designs illuminate several previously unappreciated mechanistic aspects of naturally occurring fusogens.

## Results

### Sequence redesign of the SNARE complex

To design synthetic protein-based fusion machinery, we began from the structure of the human neuronal SNARE (hnSNARE) complex. Prior to fusion, v-SNARE and t-SNARE proteins reside in opposing membranes and engage through progressive N- to C-terminal “zippering” that draws the membranes into close apposition and drives bilayer merger. This refolding reaction culminates in a highly stable postfusion complex comprising the v-SNARE VAMP2 and the t-SNAREs syntaxin-1A (Syn1A) and SNAP25 assembled into a parallel four-helix bundle (**Fig. 1A**). We hypothesized membrane fusion could be driven by engineered proteins whose lowest-energy state is a similar four-helix architecture. To enable assessment of fusion activity, we established a quantitative cell-cell fusion assay inspired by the flipped SNARE system (*19*) and viral fusogen reporter assays (*20*). Flipped versions of the v-SNARE (fVAMP2) and t-SNAREs (fSyn1A/fSNAP25) were expressed as type I membrane proteins on separate populations of HEK293T cells that also expressed T7 polymerase (T7pol) or harbored NanoLuciferase (Nluc) under the control of a T7 promoter (“v-cells” and “t-cells,” respectively) (**Fig. 1B**). Nluc reporter expression was observed specifically when v- and t-cells were mixed, reaching a level roughly 10-fold lower than when positive control cells expressing T7pol and the human fusogen hSyncytin-1 (*21*) were mixed with reporter cells (**Fig. S1A**). v-cells and t-cells also each expressed a distinct cytoplasmic fluorescent protein, allowing fusion to be visualized as double-positive syncytia (**Fig. S1B**). Fusion efficiency correlated with fSNARE expression levels, indicating that the assay quantitatively measures fusogen activity (**Fig. S1C**). Meanwhile, co-expression of fVAMP2 with flipped t-SNAREs in the same cells (*cis* expression) abolished fusion with t-cells due to *cis*-SNARE complex formation that precluded *trans*-SNARE pairing, confirming the specificity of the assay (**Fig. S1D**).

**Figure 1.**
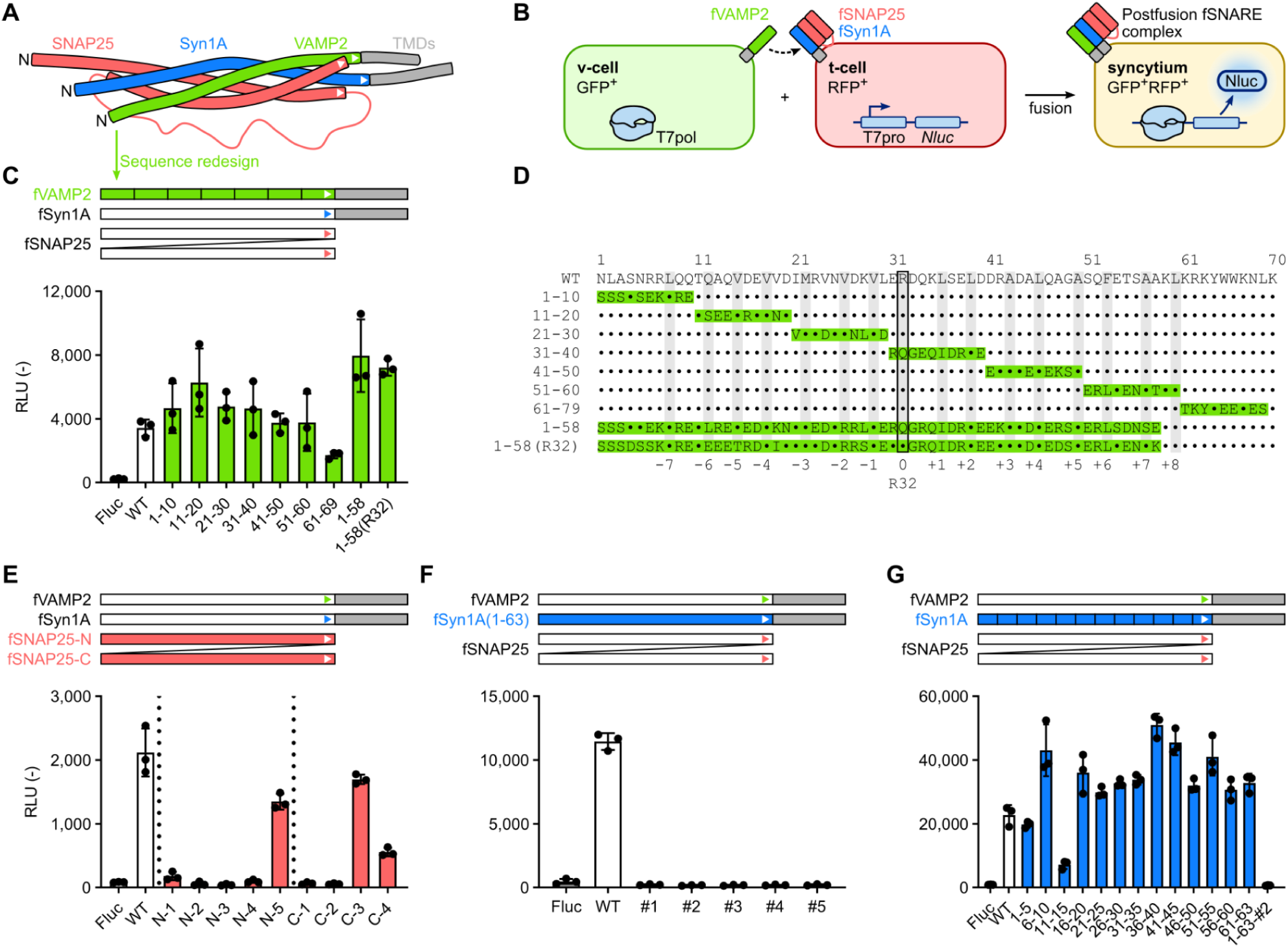
Sequence redesign of human neuronal SNARE complex. (**A**) Simplified structural schematic of the postfusion hnSNARE complex (PDB ID: 1SFC). The N-terminal end of each subunit is indicated by “N” and the directionality of each helix in the parallel four-helix bundle is indicated by a white arrowhead near its C-terminal end. TMDs, transmembrane domains. (**B**) Schematic of the cell-cell fusion assay. HEK293T cells expressing flipped v-SNARE and t-SNARE are designated as “v-cell” and “t-cell,” respectively. Cell-cell fusion efficiency was evaluated by either fluorescence microscopy or luciferase assay. (**C**) *Top*, Further simplified schematic of the postfusion SNARE complex, indicating ProteinMPNN-redesigned segments (color) and wild-type segments (white and gray). *Bottom*, Membrane fusion activity of redesigned fVAMP2 variants. Luminescence is expressed as relative luminescence units (RLU). WT fVAMP2 (white) was used as a benchmark and firefly luciferase (Fluc) was transfected instead of the fusogens as a negative control. (**D**) Sequence alignment of the coiled-coil domains of wild-type (WT) and redesigned fVAMP2 proteins. The redesigned segment in each construct is highlighted in green. The identities of redesigned residues are provided, while WT residues are represented as dots. Gray columns indicate positions at the inner core of each layer of the coiled-coil, and the ionic layer at position 32 is outlined in black. (**E**) *Top*, Schematic of sequence redesign of fSNAP25 by ProteinMPNN, colored as in **C**. Each fSNAP25 coiled-coil domain was redesigned independently. *Bottom*, Membrane fusion activity of redesigned fSNAP25 variants measured in the cell-cell fusion assay. (**F**) *Top*, Schematic of sequence redesign of fSyn1A by ProteinMPNN. *Bottom*, Membrane fusion activity of redesigned fSyn1A(1–63) variants measured in the cell-cell fusion assay. (**G**) *Top*, Schematic of redesign of fSyn1A in 5-aa segments by ProteinMPNN. *Bottom,* Membrane fusion activity of redesigned fSyn1A variants measured in the cell-cell fusion assay.

To begin designing synthetic fusogens, we used ProteinMPNN (*22*) to redesign the amino acid sequences of fVAMP2, fSyn1A, and fSNAP25 without altering their backbone structures (the amino acid sequences of all novel proteins used in this study are provided in **Supplementary Table 2**). The transmembrane domains (TMDs) of fVAMP2 and fSyn1A, as well as the juxtamembrane domain (JMD) of fSyn1A, were left unaltered because these domains are critical for fusion activity in the hnSNARE complex (*23–25*). Redesign of the fVAMP2 coiled-coil domain (residues 1–58), either in 10-residue segments or as a whole, had little effect on fusion efficiency (**Fig. 1C and 1D**). Notably, the fully redesigned variant fVAMP2-MPNN1-58 exhibited up to two-fold higher activity than native fVAMP2, despite the mutation of nearly all solvent-exposed residues and several of the hydrophobic residues in the coiled-coil core. Substitution of the conserved Arg32, which forms the ionic layer in the native hnSNARE complex (*26*), did not reduce activity. By contrast, redesigning residues 61-69 in the JMD impaired fusion, confirming the critical role of this region (*27*).

For fSNAP25, which contains two coiled-coil domains, we independently redesigned each domain while maintaining the sequence of the other and the long intervening linker (**Fig. S2A and S2B**). Several redesigned variants retained substantial activity, demonstrating that either the N- or C-terminal coiled-coil domain can be redesigned (**Fig. 1E**). By contrast, redesign of the single fSyn1A coiled-coil domain eliminated activity (**Fig. 1F and S2C**). However, redesign of fSyn1A in five-residue segments yielded variants with slightly enhanced activity except for fSyn1A(11–15), which showed a marked loss of function (**Fig. 1G and S2C**).

To further examine the sequence-dependence of fusion activity within the TMD, JMD, and N-terminal regions of SNARE components, we designed additional mutants and evaluated their activity in the cell-cell fusion assay. The TMD sequences of both fVAMP2 and fSyn1A proved dispensable, as replacement with non-native TMDs still supported robust membrane fusion, suggesting that the TMD functions primarily as a membrane anchor rather than directly contributing to fusogenic activity (**Fig. S3A**). In fVAMP2, deletion of the unstructured N-terminal region did not affect fusion, indicating that this segment is not required for activity (**Fig. S3B**). By contrast, replacing the JMD of fVAMP2 with nine consecutive alanines (fVAMP2-JMD-A9) completely abolished fusion, consistent with previous findings (*27*), again confirming the essential role of the JMD (**Fig. S3C**). Furthermore, substitution of the JMD with non-native sequences such as poly-arginine, poly-lysine, or TatP59W (*27*) markedly reduced fusion activity, indicating that the polybasic character of the JMD alone is insufficient for fusogenic function. Notably, the fusion-deficient fVAMP2-JMD-A9 mutant inhibited cell-cell fusion when co-expressed with t-SNAREs *in cis*, suggesting that the A9 mutations do not impair t-SNARE binding, but instead prevent the initiation of membrane destabilization and subsequent fusion (**Fig. S3D**).

We next generated N-terminally truncated variants of individual SNARE components. Deletion of up to 20 residues from the N terminus of fSNAP25 was tolerated, indicating that this region is not essential for coiled-coil assembly (**Fig. S3E**). By contrast, N-terminal truncations of fVAMP2 and fSyn1A generally reduced fusion activity, although the fSyn1A-ΔN5 mutant unexpectedly exhibited higher fusion activity than the wild-type (**Fig. S3E and S3F**). Together, these sequence redesign and truncation studies identified three architectural elements necessary and sufficient for functional redesigned SNARE complexes: the SNARE domain for coiled-coil formation, the fVAMP2 JMD for membrane destabilization, and TMDs to anchor the complex in opposing membranes. Among these elements, only the JMD and the Syn1A SNARE domain displayed stringent sequence requirements for efficient fusion, suggesting that it is the structure and stability of the four-helix bundle, rather than specific sequences, that are the hallmark of functional SNARE-like fusogens.

### Construction of single-chain t-SNAREs and two-component fusogens

The parallel four-helix bundle formed by the hnSNARE complex requires connection of the N- and C-terminal helices of SNAP25 through a long, intrinsically disordered linker (**Fig. 1A**). Our sequence redesign results, however, suggested that alternative architectures might be compatible with fusion activity. An antiparallel arrangement of the SNAP25 helices could simplify the overall topology by allowing them to be connected by a shorter, more compact linker. To test this possibility, we independently reversed the amino acid sequences of the two α helices of fSNAP25, removed the native linker, and reconnected the helices with a minimal linker. The resulting structures were modeled using ColabFold and used as inputs to redesign the sequences of the reversed helices with ProteinMPNN. Evaluation in the cell-cell fusion assay revealed weak but reproducible fusion activity when the originally N-terminal coiled-coil domain of fSNAP25 was flipped to the C terminus in an antiparallel orientation (**Fig. 2A**).

**Figure 2.**
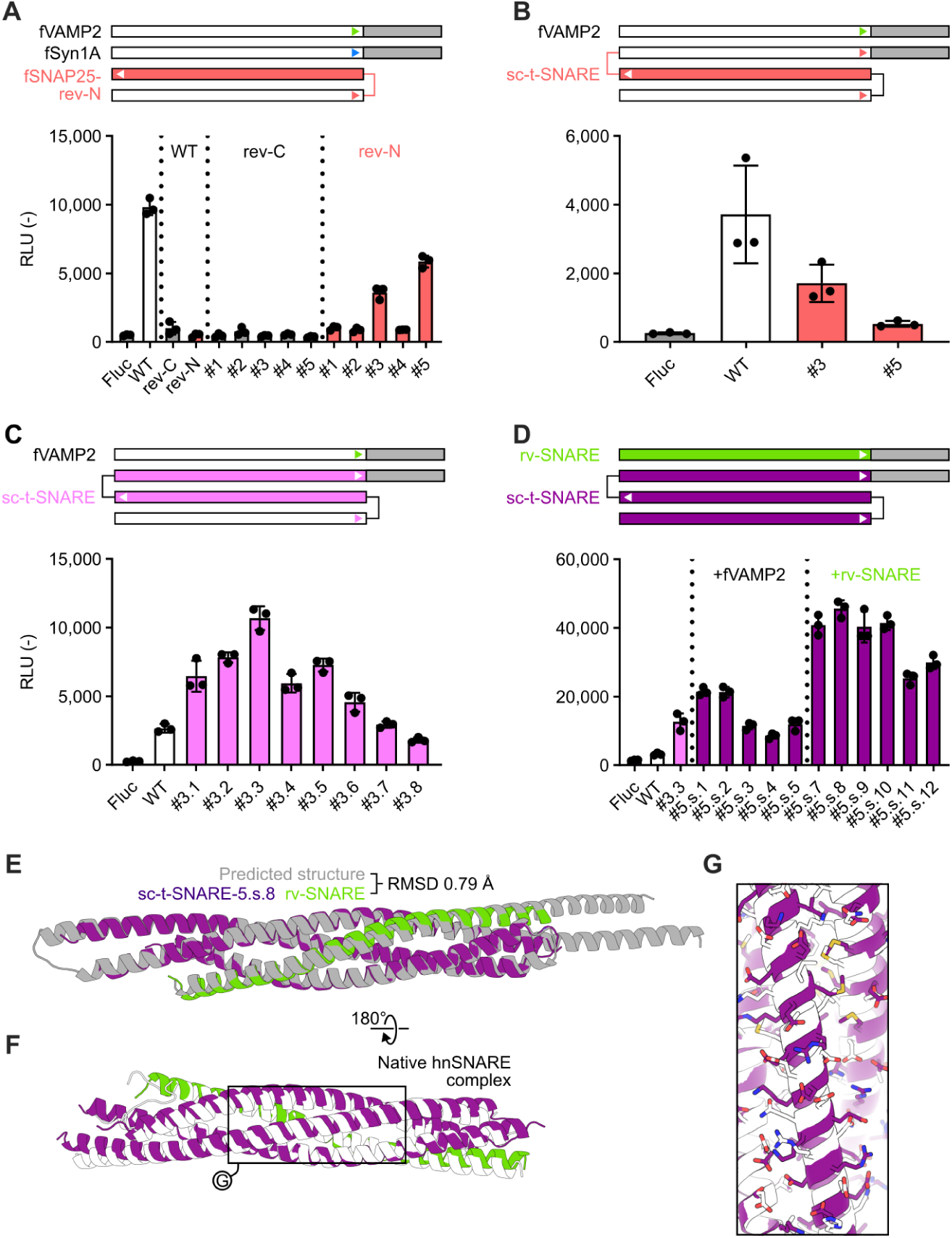
Design of single-chain t-SNAREs (sc-t-SNAREs). (**A**) *Top*, In fSNAP25-rev-N, the originally N-terminal helix of fSNAP25 was reversed, redesigned (salmon), and fused to the other SNAP25 helix via a short linker. Analogous fSNAP25-rev-C designs were also created in which the same alterations were applied to the originally C-terminal helix. *Bottom*, Membrane fusion activity of redesigned fVAMP2 variants. Luminescence is expressed as relative luminescence units (RLU). WT fVAMP2/fSyn1A/fSNAP25 (white) was used as a benchmark and firefly luciferase (Fluc) was transfected instead of the fusogens as a negative control. (**B**) *Top*, The C termini of functional fSNAP25-rev-N constructs were fused to the N terminus of fSyn1A via a short linker (salmon) to make single-chain t-SNAREs (sc-t-SNAREs). *Bottom*, Membrane fusion activity of sc-t-SNAREs, plotted as in **A**. (**C**) *Top*, Schematic of sequence redesign of sc-t-SNARE #3 by ProteinMPNN (pink). *Bottom*, Membrane fusion activity of redesigned sc-t-SNAREs. (**D**) *Top*, all coiled-coil domains in sc-t-SNARE were redesigned by ProteinMPNN (plum). *Bottom*, Membrane fusion activity of fully redesigned sc-t-SNAREs combined with either native fVAMP2 or the redesigned rv-SNARE. The WT flipped hnSNARE and fVAMP2/sc-t-SNARE-#3.3 were used as benchmarks. (**E**) Superposition of the predicted (gray) and crystal (green and plum) structures of the rv-SNARE/sc-t-SNARE-5.s.8 coiled-coil region (PDB ID: 9Y5G). (**F**) Superposition of the crystal structures of the hnSNARE complex (PDB ID: 1SFC; white) and rv-SNARE/sc-t-SNARE-5.s.8 (green and plum). (**G**) Details of the aligned structures in **F**, highlighting the register shift of the antiparallel helix in sc-t-SNARE-5.s.8 compared to the native SNAP25.

Building on this configuration, we linked the C terminus of the rearranged fSNAP25 to the N terminus of fSyn1A with a short linker, generating a single-chain t-SNARE (sc-t-SNARE). Initial sc-t-SNARE constructs generated using the two functional fSNAP25-rev-N sequences showed very weak activity (**Fig. 2B**). However, subsequent redesign of the reversed fSNAP25- and Syn1A-derived coiled-coil domains yielded variants with higher activity than the parent flipped SNARE complex (**Fig. 2C**). This result was surprising given the poor tolerance of the Syn1A SNARE domain to sequence redesign in the parent flipped SNARE context (**Fig. 1F**). Encouraged by this finding, we redesigned all helices of both the v-SNARE (“rv-SNARE”) and the sc-t-SNARE to generate two-component fusogens with entirely redesigned coiled-coil domains. These two-component fusogens exhibited exceptionally high fusion activity (**Fig. 2D**), indicating that sequence optimization of the postfusion rv-SNARE/sc-t-SNARE complex using ProteinMPNN directly improved fusion activity.

To assess design accuracy, we determined the crystal structure of the rv-SNARE/sc-t-SNARE-5.s.8 complex, which had the highest fusion activity among all constructs. The structure was determined at 2.02 Å resolution and closely matched the predicted AlphaFold2 model, with a Cα RMSD of 0.79 Å over 282 atoms and excellent agreement in both backbone and side chain conformations (**Fig. 2E** and **Supplementary Table 3**). The structure thus validated the accuracy of our machine learning-guided approach to the computational design of fusogens. In the native hnSNARE complex, a conserved ionic layer formed by Arg32 of VAMP2 and three glutamine residues from the t-SNAREs is located at the center of the coiled-coil (**Fig. S4A**). This ionic layer was absent in the synthetic fusogen, yet fusion activity was markedly enhanced (**Fig. S4B**), consistent with previous studies showing that the canonical ionic layer is not required for single-turnover fusion (*28*, *29*). Structural comparison further revealed a register shift in the α helix corresponding to the SNAP25-rev-N segment, consistent with its engineered antiparallel orientation (**Fig. 2F and 2G**).

We next explored the minimal requirements for functional two-component fusogens by evaluating dozens of designed sc-t-SNARE variants. Sequence redesign of the Syn1A region, including the JMD, produced variants that retained substantial activity, implying that the Syn1A JMD is largely dispensable (**Fig. S5A**). Consistent with earlier findings (**Fig. S3E and S3F**), truncation of 20 residues from the C terminus of the reversed fSNAP25-N helix combined with full redesign of the sc-t-SNARE coiled-coil region generated smaller variants with modest activity (**Fig. S5B**). Finally, replacing the wild-type Syn1A-derived TMD of sc-t-SNARE-5.s.8 with non-native TMDs reduced fusion efficiency (**Fig. S5C**), indicating that fusion activity is more influenced by the TMD sequence in engineered SNAREs than in the wild-type flipped hnSNARE (**Fig. S3A**).

### Generation of new fusogen backbones by RFdiffusion

Having established that a stable postfusion helix bundle is the primary structural requirement for synthetic SNARE-like fusogens, we reasoned that novel fusogen backbones could be generated computationally. We used the generative diffusion-based model RFdiffusion (*30*) to create new backbone structures based on the structure of our most active synthetic SNARE complex, rv-SNARE/sc-t-SNARE-5.s.8, followed by amino acid sequence design of the diffused regions with ProteinMPNN.

We first applied partial diffusion to the SNAP25-derived region of the sc-t-SNARE, selecting 30 novel designs for experimental characterization (**Fig. 3A**). In cell-cell fusion assays with rv-SNARE, 6 designs retained more than 50% of the parental sc-t-SNARE-5.s.8 activity, and 13 exhibited measurable activity. We next extended partial diffusion to both the SNAP25-and Syn1A-derived regions of the sc-t-SNARE (excluding the JMD and TMD), which yielded designs with more divergent backbone structures than those generated by SNAP25 diffusion alone (**Fig. 3B**). Among 15 designs tested, four showed measurable activity, and one (diff-#4.2) achieved fusion activity indistinguishable from the parental construct despite their sc-t-SNAREs sharing only 36.6% sequence identity, much of which is in the identical TMD. To quantify structural divergence introduced by partial diffusion, we compared predicted structures of designs generated using two-helix (2H) or three-helix (3H) diffusion protocols against the parental input structure. As shown in **Fig. 3C**, 3H designs exhibited significantly higher Cα RMSD values than 2H designs, indicating greater structural diversity relative to the starting template. Active and inactive designs were distributed across the RMSD range for both 2H and 3H sets, and no clear correlation was observed between RMSD and fusion activity (**Fig. 3C**), suggesting that increased backbone divergence alone does not predict functional outcome. Although partial diffusion of the sc-t-SNARE was thus tolerated, partial diffusion of rv-SNARE consistently failed to produce functional fusogens.

**Figure 3.**
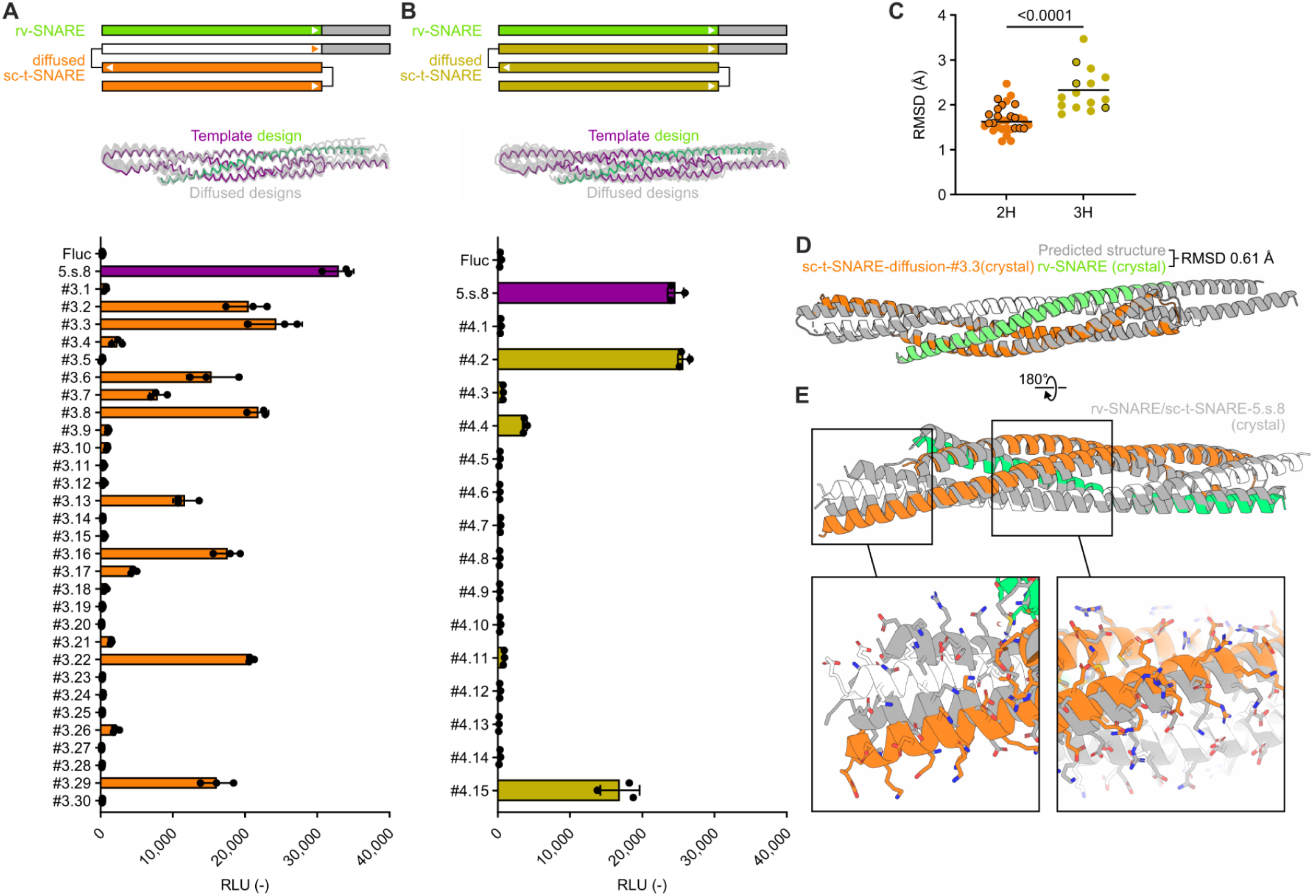
Generating synthetic fusogens with new backbones. **(A)** *Top*, Schematic of sc-t-SNARE designs in which only the SNAP25 portion (orange) was partially diffused. *Middle*, Superposition of predicted backbone structures of the template and 30 diffused designs. *Bottom*, Membrane fusion activity of diffused variants #3.1 to #3.30. Luminescence is expressed as relative luminescence units (RLU). rv-SNARE/sc-t-SNARE-5.s.8 (plum) was used as a benchmark and firefly luciferase (Fluc) was transfected instead of the fusogens as a negative control. **(B)** *Top*, Schematic of sc-t-SNARE designs in which the entire coiled-coil domain of the sc-t-SNARE was partially diffused (gold). *Middle*, Superposition of predicted backbone structures of the template and 15 designs selected for experimental characterization. *Bottom*, Membrane fusion activity of diffused variants, plotted as in **A**. (**C**) Cα RMSD values of diffused designs relative to the corresponding input template (rv-SNARE/sc-t-SNARE-5.s.8). Designs were generated by partial diffusion of either two helices (2H, orange) or three helices (3H, gold). Designs with measurable fusion activity in the cell-cell fusion assays in panels **A** and **B** are shown as outlined points. Statistical significance between 2H and 3H distributions was assessed by Welch’s t-test. (**D**) Superposition of the predicted structure (gray) and the crystal structure of the complex (PDB ID: 9Y5H) composed of rv-SNARE (green) and sc-t-SNARE-diff-#3.3 (white and orange). (**E**) Comparison of the crystal structures of the parental rv-SNARE/sc-t-SNARE-5.s.8 complex (gray, Fig. 2) and the partially diffused complex rv-SNARE/sc-t-SNARE-diff-#3.3 (white, orange, and green). Insets are close-ups of the complex with side chains shown.

To assess design accuracy, we solved the crystal structure of an active partially diffused design, rv-SNARE with sc-t-SNARE-diff-#3.3, at 2.52 Å resolution (**Fig. 3D** and **Supplementary Table 3**). The structure closely matched the AlphaFold2 (AF2)-predicted design model, yielding a Cα RMSD of 0.605 Å over 266 atoms, while differing modestly from the parental rv-SNARE/sc-t-SNARE-5.s.8 complex (Cα RMSD of 1.37 Å over 262 atoms; **Fig. 3E**). As expected, much of the deviation from the parental complex was concentrated in the SNAP25-derived coiled-coil domain, whereas the rv-SNARE remained relatively unchanged.

### Characterization of two-component fusogens

The kinetics of cell-cell fusion in the experiments above remained unclear because the reporter gene-based assay provided only endpoint measurements after 18–24 h of incubation. To address this, we employed an alternative assay based on reconstitution of split NanoLuc (Nluc), in which cell-cell fusion immediately reconstitutes active Nluc, enabling real-time monitoring of fusion kinetics (*31*). The synthetic fusogen with the highest endpoint activity (rv-SNARE/sc-t-SNARE-5.s.8) induced detectable fusion within 10 min, with luminescence reaching a plateau after 40–50 min (**Fig. 4A**). By contrast, the native hnSNARE complex displayed slower kinetics, with fusion detectable only after ∼20 min, and reached a considerably lower maximum signal. Fluorescence imaging further showed that rv-SNARE with sc-t-SNARE-5.s.8 produced multinucleated syncytia more efficiently than the native flipped hnSNARE complex (**Fig. 4B**), illustrating the superior fusion activity of the synthetic designs.

**Figure 4.**
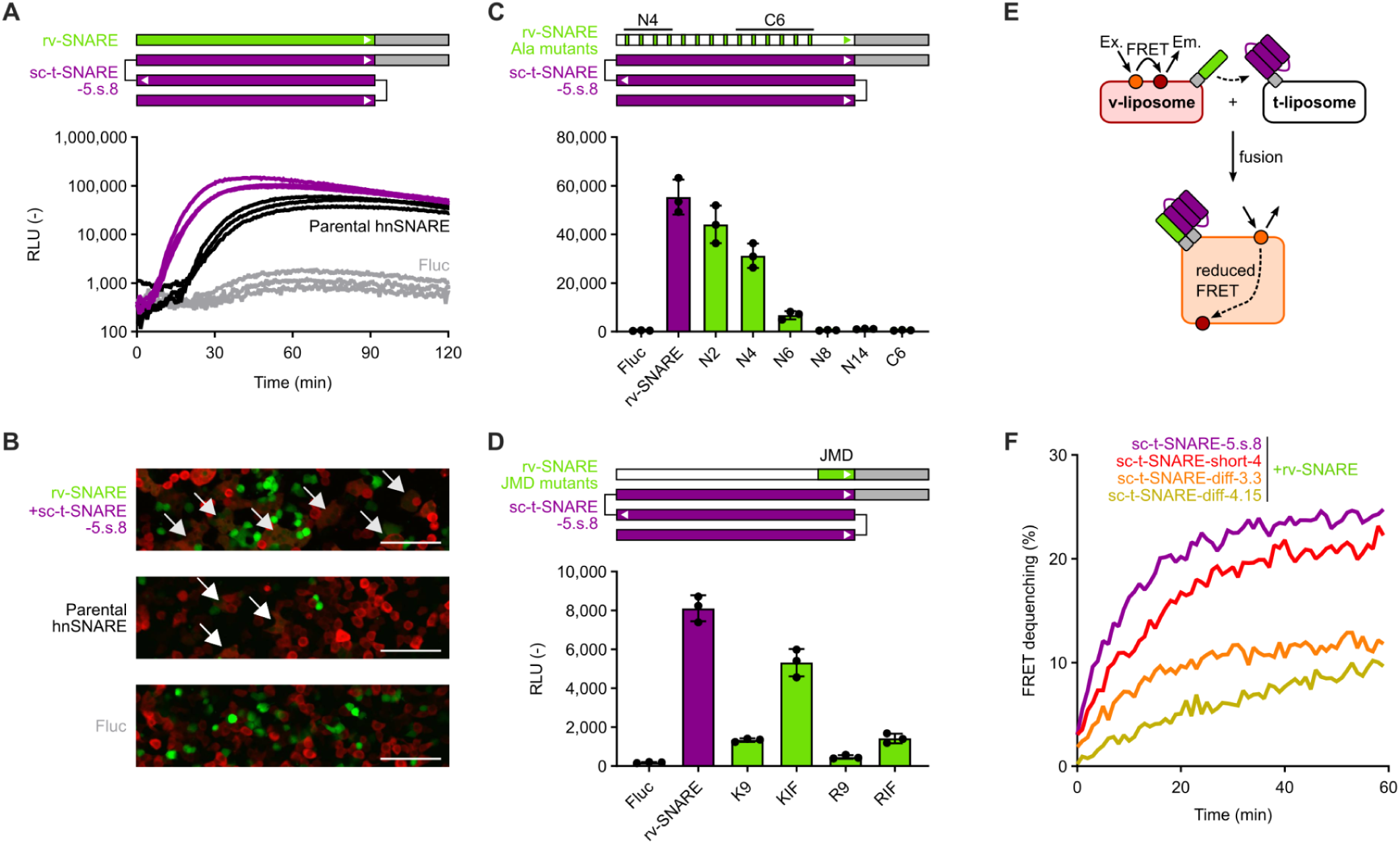
Characterization of two-component fusogens. (**A**) Kinetics of cell-cell fusion measured by split Nluc complementation. The experiment was performed in triplicate, and all data are presented. (**B**) Confocal microscope images of HEK293T cells after 2 h of cell-cell fusion. Cells expressing the v-SNARE and t-SNARE were labeled with mNeonGreen (green) and DsRed (red), respectively. Arrows indicate multinucleated cells formed by cell-cell fusion. Scale bars, 100 µm. (**C**) Membrane fusion activity of rv-SNARE variants with multiple mutations of hydrophobic residues at the interface to alanine. (**D**) Membrane fusion activity of rv-SNARE variants with mutant JMD sequences (**Supplementary Table 2**). (**E**) Schematic of lipid mixing assay to measure synthetic SNARE-mediated liposomal fusion. Fluorescently labeled liposomes displaying rv-SNARE (v-liposome) were mixed with non-labeled liposomes displaying sc-t-SNARE (t-liposome). (**F**) Fusion activity, measured as donor fluorescence emission, of rv-SNARE with sc-t-SNARE variants in the lipid mixing assay. Representative results are shown (n=2 for each experimental group).

We next examined why the synthetic fusogens outperformed the parent flipped hnSNARE complex. One possibility was enhanced expression in mammalian cells, as ProteinMPNN-redesigned proteins often express at higher levels (*32*). We tested this hypothesis by measuring the expression level of synthetic fusogens in HEK293T cells by genetically fusing them to the luminescence tag HiBiT (*33*). Indeed, rv-SNARE and t-SNAREs derived from sc-t-SNARE-5.s.8 were expressed at higher levels than their native counterparts (**Fig. S6A and S6B**). However, expression alone did not fully explain activity differences: sc-t-SNARE-5.s.8 and sc-t-SNARE-#3.3 were expressed at comparable levels, yet 5.s.8 exhibited markedly higher fusion activity (**Fig. S6B and S6C**). Similarly, sc-t-SNARE-#5 was expressed at only two-fold lower levels than 5.s.8 but displayed >30-fold lower activity. These results suggest that expression may contribute to, but is not solely responsible for, the enhanced activity of the synthetic fusogens.

We then focused on structural determinants. Because the postfusion SNARE complex is stabilized largely by hydrophobic contacts, we hypothesized that interface hydrophobicity governs complex affinity and fusogenicity. Fourteen hydrophobic residues in rv-SNARE are present at the interface with the t-SNARE (**Fig. S6D**). Alanine substitutions at individual positions minimally affected activity, except at the C-terminal 14th residue (**Fig. S6E**). However, progressive accumulation of alanine substitutions caused a marked loss of activity (**Fig. 4C and S6D**), indicating that robust hydrophobic interactions are essential for functionality. Interestingly, fusion-deficient alanine mutants of rv-SNARE still bound to sc-t-SNARE-5.s.8 *in cis* and completely blocked cell-cell fusion (**Fig. S6F**), suggesting that weak interactions are insufficient to drive fusion but still form stable inhibitory complexes. Together, these data imply that there might be a threshold of affinity that determines functionality as a fusogen.

Finally, we re-examined the role of the JMD in the context of our two-component fusogens. In native fVAMP2, the polybasic JMD is critical for fusion (**Fig. S3C**), likely serving to destabilize the embedded and opposed membranes (*27*). “KIF” and “RIF” rv-SNARE variants, in which positively charged residues in the native JMD were replaced with Lys or Arg, respectively, while hydrophobic and aromatic residues were replaced with Ile and Phe, showed higher activity than poly-Lys (K9) or poly-Arg (R9) mutants, indicating that a combination of positive charge and hydrophobicity is required (**Fig. 4D**). The complete loss of activity in KIF and RIF shuffle variants demonstrated that residue positioning is also critical (**Fig. S6G**). Furthermore, KIF and KIW variants outperformed KIY and KIS, highlighting the importance of hydrophobic aromatic side chains in the JMD (**Fig. S6H**).

We also examined the orthogonality of the designed fusogens using two v-SNARE variants and three sc-t-SNARE variants. rv-SNARE exhibited robust compatibility with all sc-t-SNARE variants, whereas fVAMP2-WT supported only weak fusion activity overall and showed particularly low activity when paired with sc-t-SNARE-short-4 (**Fig. S7**). These findings demonstrate that orthogonal v-SNARE/sc-t-SNARE pairs can be rationally designed.

### In vitro reconstitution of synthetic fusogens

We next tested whether the synthetic fusogens were sufficient to drive membrane fusion in synthetic liposomes using a lipid-mixing assay based on Förster resonance energy transfer (FRET) between the lipophilic dyes DiI and DiD (*31*) (**Fig. 4E**). Liposomes composed of DOPC and cholesterol (± DiI/DiD) were prepared, and recombinant rv-SNARE and sc-t-SNARE variants were reconstituted into the labeled and unlabeled liposomes, respectively (**Fig. S8A**). Mixing rv-SNARE–containing liposomes with sc-t-SNARE–containing liposomes produced a FRET change consistent with lipid mixing within minutes (**Fig. 4F**), indicating that the synthetic fusogens can drive membrane fusion without exogenous triggers, unlike rapid, Ca²⁺-triggered neuronal fusion, which requires accessory proteins (*35*). An excess of soluble rv-SNARE competitively inhibited fusion, confirming that lipid mixing depended on specific interactions of membrane-anchored rv-SNAREs with sc-t-SNAREs and the formation of a trans-SNARE complex (**Fig. S8B to S8D**). Nine consecutive alanine substitutions in the rv-SNARE JMD abolished fusion (**Fig. S8E to S8G**), consistent with the cell-cell fusion results for the JMD-A9 mutant (**Fig. S3D**). Overall, these lipid-mixing data closely mirror the results of cell-cell fusion assays, indicating that the synthetic fusogens operate robustly across the membrane curvatures and lipid compositions tested.

### Chemically inducible fusogens

We next engineered chemically inducible fusogens by integrating into the synthetic fusogens the rapamycin-dependent FKBP-FRB heterodimer (*36*). FKBP and FRB were fused to the N termini of v-SNAREs and sc-t-SNAREs, respectively, and fusion activity was assessed using cell-cell fusion assays in the presence or absence of rapamycin (**Fig. 5A**). In the absence of rapamycin, the FKBP and FRB domains almost completely suppressed basal fusion activity of wild-type fVAMP2-based constructs, whereas some basal activity remained when using FKBP-fused rv-SNARE (**Fig. S9A**). In both cases, the addition of 50 nM rapamycin resulted in cell-cell fusion, the extent of which depended on the identity of the sc-t-SNARE used. These results suggest that N-terminal steric hindrance inhibited the initiation of *trans*-SNARE complex formation, but that this inhibition could be overcome by the presumably higher-affinity rv-SNARE/sc-t-SNARE interaction or, controllably, by chemically induced dimerization.

**Figure 5.**
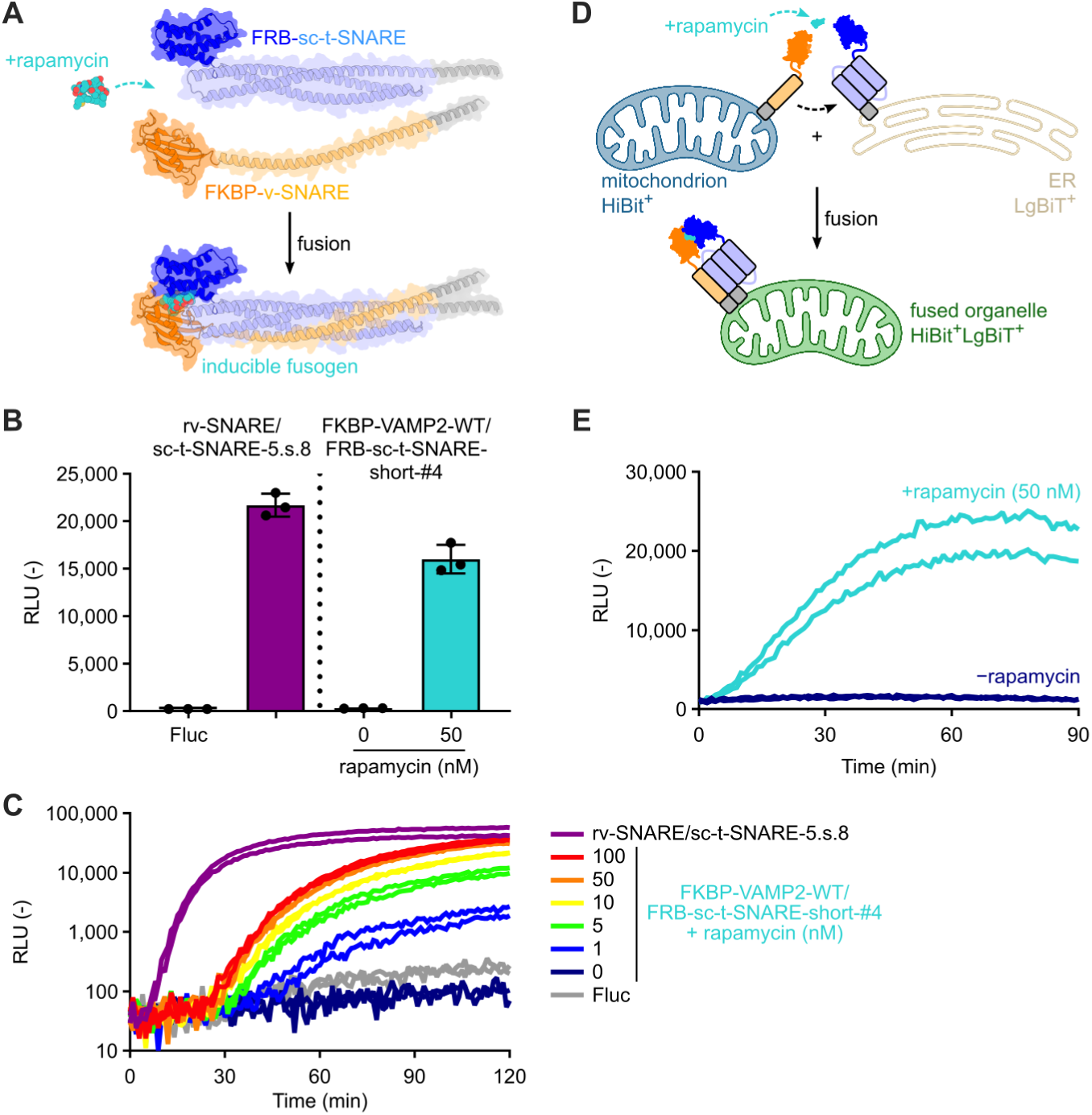
Cell-cell fusion and ER-mitochondrion fusion by chemically inducible fusogens. (**A**) Schematic of a rapamycin-inducible fusogen. In the presence of rapamycin, heterodimerization of the FK506-binding protein (FKBP) and FKBP–rapamycin-binding (FRB) domains (PDB ID: 3FAP) at the N terminus of synthetic fusogens initiates complex formation and induces fusion. (**B**) Rapamycin-dependent cell-cell fusion by the inducible fusogen FKBP-fVAMP2-WT/FRB-sc-t-SNARE-short-#4, measured by T7-Nluc reporter assay. (**C**) Kinetics of cell-cell fusion by the inducible fusogen, measured by split Nluc complementation assay. The non-inducible fusogen rv-SNARE/sc-t-SNARE-5.s.8 was used as a reference. The experiment was performed in duplicate, and all data are presented. (**D**) Schematic of ER-mitochondrion fusion assay. Rapamycin-dependent ER-mitochondrion fusion reconstitutes active Nluc from MT-HiBiT and ER-LgBiT. (**E**) Kinetics of Nluc activity in ER-MT fusion assay. The experiment was performed in duplicate, and all data are presented.

Pairing FKBP-fVAMP2-WT with FRB-sc-t-SNARE-short-#4 resulted in switch-like behavior in which fusion was both efficient and completely dependent on the presence of 50 nM rapamycin (**Fig. 5B**). Notably, the parental, non-inducible form of this combination showed almost no fusion activity (**Fig. S7**), indicating that rapamycin-induced dimerization can rescue inefficient v-SNARE/sc-t-SNARE pairs. Kinetic analysis revealed that inducible fusion depended on rapamycin concentration and proceeded more slowly than non-inducible fusion. Although these fusogens incorporate different coiled-coil backbones and are therefore not directly comparable, the non-inducible rv-SNARE/sc-t-SNARE-5.s.8 (used as a benchmark due to its high efficiency) initiated fusion within 10 min, whereas the inducible FKBP-fVAMP2-WT/FRB-sc-t-SNARE-short-#4 required ∼30 min after mixing v- and t-cells (**Fig. 5C**). Finally, segment-swapping experiments between wild-type fVAMP2 and rv-SNARE demonstrated that the N-terminal region of fVAMP2, particularly residues 11–20, contributes to inducibility by conferring low basal activity combined with strong rapamycin responsiveness, although the underlying mechanism remains unclear (**Fig. S9B and S9C**).

### Organelle fusion using inducible synthetic fusogens

We hypothesized that, like naturally occurring SNARE complexes, our synthetic fusogens may be able to fuse intracellular membranes. As a proof of concept, we generated constructs intended to target the fusogens to the cytoplasmic side of the endoplasmic reticulum (ER) and outer mitochondrial membranes to drive organelle-organelle fusion. An mNeonGreen (mNG)-tagged variant of sc-t-SNARE-5.s.8 bearing the wild-type Syn1A TMD predominantly accumulated in the ER, while mNG-rv-SNARE bearing the wild-type VAMP2 TMD localized to the plasma membrane as well as the ER (**Figs. S10A and S10B**). By contrast, the TMD and tail of Fis1 (*37*) directed both synthetic fusogens to the mitochondrial outer membrane (MOM) (**Fig. S10A**), while the C-terminal ER retention signal KDEL (*38*) strongly localized them to the ER.

To assess organelle-organelle fusion, we developed an in vitro content mixing assay (**Fig. 5D**). MOM-localized FKBP-fVAMP2-WT-Fis1 and ER-localized FRB-sc-t-SNARE-short-#4-KDEL (**Figs. S11A and S11B**) were expressed in separate cell populations, together with split Nluc probes targeted to the mitochondrial intermembrane space (fusion with LACTB protein) and ER lumen (KDEL), respectively (*38*, *39*). After cell disruption, isolated mitochondria and ER showed gradual Nluc reconstitution only when mixed in the presence of rapamycin, indicating synthetic fusogen-mediated ER-MOM fusion (**Fig. 5E**). No signal was observed when the HiBiT reporter was instead targeted to the mitochondrial matrix, consistent with fusion being restricted to the mitochondrial outer membrane and not involving inner-membrane fusion or matrix content mixing (**Fig. S11C**). Co-expression of soluble, cytoplasmic rv-SNARE in t-SNARE–expressing cells reduced luminescence to background levels, indicating that fusion required *trans*-SNARE complex formation (**Fig. S11D**). These findings provide proof of principle that synthetic fusogens can be directed to specific organelles and fuse intracellular membranes.

## Discussion

This study establishes a general framework for the computational design of protein-based membrane fusion machinery. Using machine learning-guided design tools, we generated nearly 200 functional synthetic fusogens spanning diverse sequences and architectures. This work extends beyond the original flipped SNARE system and prior synthetic fusogen studies by definitively identifying *trans*-SNARE engagement and formation of a stable postfusion coiled-coil complex as the minimal requirements for membrane fusion. Our ability to generate large numbers of active fusogens with widely varying sequences—and even altered topologies—shows that neither specific amino acid sequences nor a particular backbone architecture are required for productive fusion. With continued advances in generative protein design it should be possible to explore an even broader landscape of functional fusogen architectures.

The molecular basis for the exceptional activity of the synthetic fusogens remains to be fully resolved. Elevated expression levels may contribute in part, but ProteinMPNN-guided redesign likely optimized the interactions between the v-SNARE and t-SNARE (27), increasing the efficiency of productive trans-complex formation. In addition, consolidation of the two t-SNARE components into a single sc-t-SNARE molecule likely accelerates fusion by bypassing the multistep assembly required for the native hnSNARE complex.

In contrast to endogenous neuronal SNARE-mediated fusion—which is tightly regulated by accessory proteins that control triggering, clamping, and recycling (34)—the synthetic fusogens described here operate autonomously. By using unregulated, single-turnover fusion assays, this study isolates the activity of the core fusogenic machinery and extends to synthetic fusogens earlier observations showing that membrane fusion can proceed in the absence of the native regulatory network (*19*).

The synthetic fusogens function across multiple contexts, including cell-cell fusion, reconstituted liposomal systems, and organelle fusion between the ER and mitochondria. Their genetic encodability and modular design enable straightforward integration of additional functionality, such as chemically inducible control and subcellular targeting. This modularity suggests a clear path toward engineering fusogens that respond to defined molecular cues or act selectively on specific cells or membranes.

Several limitations should be noted. First, the fusion assays used here primarily report unregulated, single-turnover events, and the synthetic fusogens do not recapitulate the full regulatory complexity or recycling behavior of native SNARE-mediated fusion. Second, several of our cell-cell fusion assays show that the individual components of the synthetic fusogens are not fully orthogonal to native SNARE proteins. Designing bio-orthogonality, which we show is possible, will be desirable for many synthetic biology applications. Finally, although the current designs rely on two-component systems, many therapeutic applications will ultimately require reduction to a single-component fusogen. Achieving this goal presents a substantial challenge, but the design principles and modular strategies established here provide a foundation for future efforts toward single-chain, regulatable fusogens capable of targeted delivery.

Together, these findings establish computationally designed protein fusogens as a versatile and extensible class of molecular tools. By defining the minimal requirements for membrane fusion and demonstrating broad sequence and architectural tolerance, this work opens new opportunities for engineering membrane dynamics in synthetic biology and for developing programmable fusion systems for therapeutic delivery.

## Materials and Methods

### Computational methods

For protein sequence redesign, we employed ProteinMPNN (*22*), using the crystal structure of the postfusion hnSNARE complex (PDB ID: 1SFC) (*26*) or AF2-predicted structures as input. Output sequences were subjected to structure prediction using ColabFold (*40*, *41*) and sequences were selected for experimental characterization based on pLDDT scores and visual inspection using the PyMOL Molecular Graphics System versions 2.5 to 3.0 (Schrödinger).

RFdiffusion (*30*) was used for the generation of new backbones by partial diffusion, using the AF2-predicted structure of the rv-SNARE/sc-t-SNARE-5.s.8 complex as input. Certain portions of the sc-t-SNARE protein were fixed, and other regions were partially diffused. For partial diffusion, diffuser.partial_T was set to 10, 20, or 30. Amino acid sequences for the resultant backbones were designed by ProteinMPNN, and candidate designs were filtered based on the pLDDT scores of AF2-predicted structures, RMSD against input structures, and visual inspection.

### Biological materials and cell culture

Synthetic genes were obtained from Integrated DNA Technologies and cloned into the expression vector pCMV, pcDNA3.1, or pET29b by Gibson assembly (*42*) or Golden Gate cloning (*43*). DNA plasmids used in this study are listed in **Supplementary Table S1**.

HEK293T cells were obtained from ATCC and maintained in DMEM supplemented with 10% fetal bovine serum and antibiotics at 37°C with 5% CO_2_.

### Cell-cell fusion assays

The endpoint reporter gene-based cell-cell fusion assay was inspired by an existing fusion assay for viral fusogens (*20*) with slight modification. Cells expressing v-SNARE proteins co-expressed T7 polymerase and the green fluorescence protein mNG, while cells expressing t-SNARE proteins co-expressed the red fluorescence protein DsRed and harbored an Nluc gene under the T7 promoter. Synthetic SNARE-like proteins were expressed on the surface of HEK293T cells as flipped SNAREs (*19*). HEK293T cells were seeded on day 0 in 96-well plates (white color, clear bottom) at a density of 2.5×10^4^ cells per well, and transfected using PEI Max reagent (Polyscience) with a total of 100 ng plasmid DNA per well on day 1 (50 ng v-SNARE, 20 ng T7 polymerase, and 30 ng mNG for v-cells; 50 ng t-SNARE, 10 ng T7-Nluc, and 40 ng DsRed for t-cells; Fluc plasmid was used instead of SNAREs for negative controls). On day 2, the culture medium was exchanged and v-cells were resuspended in fresh medium by pipetting, added to the t-cells, and cultured overnight. On day 3, the culture medium was removed and the Nluc substrate furimazine (Promega) was added to the cells. Luminescence signal was measured by plate reader using Synergy 2, Synergy Neo2, or Synergy H1 plate readers (Agilent). Assays were performed in triplicate.

For the real-time cell-cell fusion assay, we employed the split Nluc system (*31*, *44*). HEK293T cells were seeded on day 0 in 96-well plates (white color, clear bottom) at a density of 2.5×10^4^ cells per well. Cells were transfected on day 1 using PEI Max reagent with a total of 100 ng of plasmid DNA per well (50 ng v-SNARE, 50 ng HiBiT-EGFP for v-cells; 25 or 50 ng t-SNARE, 10 ng FRB-LgBiT, 40 ng DsRed, and 25 or 0 ng Fluc [filler] for t-cells). On day 2, the culture medium was removed and 100 µL per well of HBSS (+) containing 1 µM of DrkBiT peptide was added to the transfected cells. Nluc live cell substrate (Promega) was added to the t-cells before mixing with v-cells. The v-cells were detached and resuspended in the buffer by pipetting and added to the t-cells. Nluc activity was immediately and continuously measured by plate reader for up to 2 h (in 30- or 60-s intervals) at 37°C. Assays were performed in duplicate or triplicate.

### Confocal microscopy

HEK293T cells were seeded on an 8-well glass-bottomed chamber or 96-well glass-bottomed plate and transfected with pDNA using PEI as described above. Synthetic fusogens targeting the ER or MOM were tagged with mNG or split mNG_11_ (*45*), while those targeting the mitochondrial (MT) matrix and ER were labeled with mScarlet-I (*46*). The following day, fluorescence images of live cells were captured using an FV1000D confocal microscope (Olympus). Fluorescence images were processed using the Fiji image processing software, version 2.14.0.

### Protein expression and purification

Recombinant synthetic fusogens were expressed and purified using a protocol adapted from previously reported methods for SNARE proteins (*34*). Synthetic genes encoding the fusogens, each containing an N-terminal hexahistidine tag, were cloned into the pET21 vector and transformed into *Escherichia coli* BL21(DE3) or C41(DE3) strains. Cultures (200–500 mL) were grown in Terrific Broth (TB) supplemented with the appropriate antibiotic (carbenicillin or kanamycin) at 37°C to an optical density at 600 nm (OD_600_) of 0.4–0.6. Protein expression was then induced with 0.5 mM isopropyl β-D-1-thiogalactopyranoside (IPTG), and cultures were shaken overnight at 18°C. Cells were harvested by centrifugation, washed with phosphate-buffered saline (PBS), and lysed in lysis buffer (see below) by sonication. Cell debris was removed by centrifugation at 5,000–10,000 *g*, and the clarified supernatant was applied to Ni-NTA resin. After washing with lysis buffer, bound proteins were eluted using elution buffer. Protein concentrations were determined by absorbance at 280 nm or by bicinchoninic acid (BCA) assay. For proteins containing transmembrane domains, the following buffers were used: lysis buffer consisted of 25 mM HEPES pH 7.4, 400 mM KCl, 10% (v/v) glycerol, 20 mM imidazole, 4% (v/v) Triton X-100, DNase I, and protease inhibitor cocktail (Roche). Elution buffer consisted of 25 mM HEPES pH 7.4, 400 mM KCl, 10% (v/v) glycerol, 1% *n*-octyl-β-D-glucoside (β-OG), and 500 mM imidazole. For recombinant proteins lacking transmembrane domains, purification was performed using the following buffers: lysis buffer consisted of 20 mM Tris pH 8.0, 500 mM NaCl, 30 mM imidazole, DNase I, and protease inhibitor cocktail. Elution buffer consisted of 20 mM Tris pH 8.0, 150 mM NaCl, and 500 mM imidazole.

For redesigned v-SNAREs, superfolder GFP (sfGFP) was fused to the N terminus to enhance expression levels in *E. coli* and improve protein stability during purification, as recombinant rv-SNARE lacking the sfGFP tag tended to aggregate following nickel-affinity purification. The presence of the N-terminal sfGFP tag did not significantly affect fusion activity in lipid-mixing assays.

For sc-t-SNARE constructs, fusion of a SUMO tag to the N terminus enhanced expression in *E. coli*. However, sc-t-SNARE proteins could also be purified without the SUMO tag, and SUMO-free constructs did not exhibit detectable aggregation.

### X-ray crystallography

sfGFP-tagged rv-SNARE and SUMO-tagged t-SNAREs were cleaved using tobacco etch virus (TEV) protease and separated from the tags using Ni-NTA resin. SNARE complexes were formed by mixing the proteins in a 1:1 molar ratio, and were subsequently purified using SEC. Pure fractions were collected, pooled, and concentrated prior to crystallization. Crystallization trials were set up using a Mosquito LCP crystallization robot (SPT Labtech) and imaged using UVEX microscopes and a UVEX PS-256 imaging system (JAN Scientific). Diffraction-quality crystals were obtained under the following conditions:

- rv-SNARE/sc-t-SNARE-diff-#3.3 complex: 0.08 M sodium cacodylate trihydrate pH 6.5, 20% (v/v) glycerol, 0.16 M calcium acetate hydrate, and 14.4% (w/v) polyethylene glycol (PEG) 8000;
- rv-SNARE/sc-t-SNARE-5.s.8 complex (space group P4_1_2_1_2): 0.2 M ammonium acetate pH 7.1 and 20% (w/v) PEG 3350;
- rv-SNARE/sc-t-SNARE-5.s.8 complex (space group P2_1_2_1_2_1_): 0.1 M succinic acid pH 7.0 and 15% (w/v) PEG 3350.

Diffraction data were collected at the Advanced Light Source (ALS) on beamline 2.0.1 for the rv-SNARE/sc-t-SNARE-diff-#3.3 complex and at the National Synchrotron Light Source II (NSLS-II) on beamline 17-ID-2 (FMX) for the rv-SNARE/sc-t-SNARE-5.s.8 complex. Diffraction images were indexed and integrated using XDS (*47*), and data were merged and scaled using Pointless and Aimless within the CCP4 program suite (*48*). Initial phases were obtained by molecular replacement using Phaser (*49*), with the design models for each complex used as search templates. Following molecular replacement, automated model building was performed using phenix.autobuild with rebuild-in-place disabled and simulated annealing enabled. Iterative refinement was carried out using Phenix (*50*), with manual model adjustment performed in Coot (*51*). Model quality was assessed using MolProbity (*52*). Data collection and refinement statistics are summarized in **Supplementary Table 3**. Atomic coordinates and structure factors have been deposited in the Protein Data Bank (PDB; http://www.rcsb.org) under accession codes 9Y5H, 9Y5F, and 9Y5G.

### Proteoliposomes and lipid mixing assay

Liposomes were prepared using the ethanol injection method as previously reported (*53*). Briefly, lipids dissolved in ethanol (DOPC:cholesterol, 60:40 mol/mol) were injected into SNARE reconstitution buffer (25 mM HEPES pH 7.4, 100 mM KCl, 10% glycerol, 1% β-OG) to generate a lipid-detergent micelle solution. For fluorescent labeling, the lipophilic dyes DiI and DiD were included in the lipid formulation at final molar ratios of DOPC/cholesterol/DiI/DiD = 59:40:0.5:0.5. Proteoliposomes were prepared as described previously (*34*): recombinant proteins dissolved in SNARE storage buffer (25 mM HEPES pH 7.4, 400 mM KCl, 10% glycerol, 1% β-OG) were mixed with the lipid-detergent micelles. For v-SNARE proteoliposomes, 500 µg of recombinant protein was mixed with 500 µg of lipid, whereas for sc-t-SNARE proteoliposomes, approximately 50–100 µg of recombinant protein was mixed with 5,000 µg of lipid. Fluorescently labeled liposomes contained v-SNARE proteins, whereas unlabeled liposomes contained sc-t-SNARE proteins. Proteoliposome formation was achieved by overnight dialysis against SNARE reconstitution buffer to remove detergent. Proteoliposomes were subsequently purified from free protein and lipid by OptiPrep density gradient centrifugation. The resulting proteoliposomes had diameters ranging from 100 to 300 nm, as determined by dynamic light scattering using a Zetasizer Nano ZS (Malvern Panalytical).

For lipid-mixing assays, labeled v-SNARE proteoliposomes were mixed with unlabeled sc-t-SNARE proteoliposomes at volume ratios ranging from 1:2 to 1:4 (v-liposomes:t-liposomes) in SNARE reconstitution buffer in black 96-well plates. Lipid mixing was monitored by measuring DiI fluorescence (excitation at 530 nm, emission at 560 nm) using a plate reader. Fluorescence obtained after addition of 1% Triton X-100 to labeled liposomes, which eliminates FRET by complete membrane solubilization, was defined as the maximal signal and used to normalize fusion efficiency.

### In vitro organelle fusion assay

For the detection of synthetic fusogen-mediated fusion between ER and MOM, HEK293T cells in a 12-well plate were transfected to express synthetic SNAREs and NanoBiT probes. The following day, cells were washed with PBS, resuspended in PBS, and disrupted by passing through a 30G needle 10 times. The crude lysates from v-cells (FKBP-VAMP2-WT-Fis expressed on MOM and HiBiT-tagged mitochondria marker) and t-cells (FRB-sc-t-SNARE-short-4-KDEL and ER-localized LgBiT protein) were diluted in PBS containing 1 µM-DrkBiT peptide, and mixed in a white 96-well plate with Nluc substrate. The luminescence signal was continuously monitored for up to 2 hours at 37°C.

## Acknowledgments

We thank Alex Merz and Emma Mackey at the University of Washington for technical assistance on proteoliposome preparations; Yongli Zhang at Yale University and Seiichi Koike at University of Toyama for helpful discussions; Cyrus Haas for assistance with figures Kandise VanWormer, Hernan Nunez-Ortega, and Eryn Weston for laboratory and administrative support; and Luki Goldschmidt, Patrick Vecchiato, and Bulat Faezov for building and maintaining the computing infrastructure at the Institute for Protein Design. This work was supported by the Japan Society for the Promotion of Science (JSPS) Overseas Research Fellowships (M.S.), JSPS KAKENHI (25H02269 to M.S.), Japan Science and Technology Agency PRESTO, (JPMJPR24OB to M.S.), Terumo Life Science Foundation (24-III3009 to M.S.), Astellas Foundation for Research on Metabolic Disorders (2024A2123 to M.S.), “Crossover Alliance to Create the Future with People, Intelligence and Materials” from MEXT, Japan (M.S.), Takeda Science Foundation (M.S.), the Gates Foundation (INV-043758 to N.P.K.), the National Institute for Allergy and Infectious Disease (U54AI170856 to N.P.K.), and a generous gift from the Audacious Project (N.P.K.). We thank the Advanced Light Source (ALS) beamline 2.0.1 at Lawrence Berkeley National Laboratory for X-ray crystallographic data collection. The Berkeley Center for Structural Biology is supported in part by the National Institutes of Health (NIH), National Institute of General Medical Sciences, and the Howard Hughes Medical Institute. The ALS is supported by the Director, Office of Science, Office of Basic Energy Sciences and US Department of Energy (DOE) (DE-AC02-05CH11231). We also want to thank the National Synchrotron Light Source II. The Center for Bio-Molecular Structure (CBMS) is primarily supported by the NIH-NIGMS through a Center Core P30 Grant (P30GM133893), and by the DOE Office of Biological and Environmental Research (KP1607011). NSLS-II is a U.S. DOE Office of Science User Facility operated under Contract No. DE-SC0012704. This publication resulted from data collected using beamtime obtained through NECAT BAG proposal #311950. Some of the plasmids in this study were gifts from Dorus Gadella through Addgene (#98818 and #137805). Some of the illustrations in this manuscript were created by BioRender. The authors acknowledge the use of ChatGPT (OpenAI) for assistance in improving language and readability. All scientific content was generated, verified, and approved by the authors.

## Supplementary Materials

**Fig. S1.**
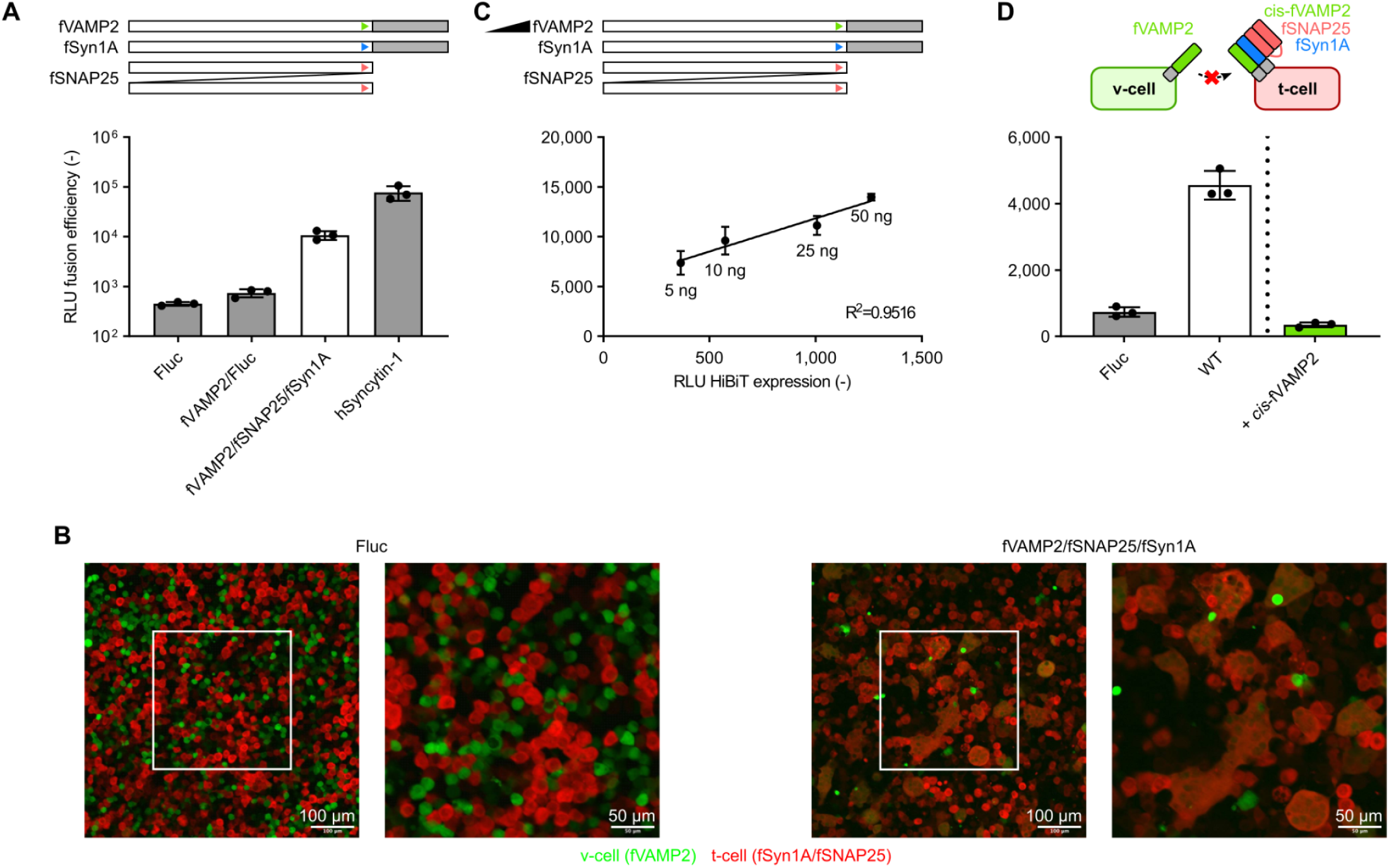
Characterization of flipped hnSNARE cell-cell fusion assay. (**A**) Cell-cell fusion assay using flipped human neuronal SNARE (hnSNARE) proteins and human Syncytin-1 (hSyncytin-1). Flipped VAMP2 (fVAMP2) was expressed in v-cells and flipped SNAP25 and Syn1A (fSNAP25 and fSyn1A) were expressed in t-cells. As a benchmark, hSyncytin-1 was expressed in t-cells and mixed with v-cells expressing no v-SNARE. Negative controls included mixing v-cells with t-cells lacking t-SNAREs (fVAMP2/Fluc) and cells lacking flipped SNAREs but expressing firefly luciferase (Fluc)Fluc. (**B**) Confocal microscopy images of cells after 24 h of cell-cell fusion by flipped hnSNAREs. The v-cells expressed cytoplasmic mNeonGreen while the t-cells expressed cytoplasmic DsRed. Cells lacking flipped SNAREs but expressing Fluc were used as a negative control. Right panels are magnified images from the boxed areas of left panels; differences in fluorescence intensity for certain cells are the result of altered focus at higher magnification. (**C**) Dependence of cell-cell fusion efficiency on fVAMP2 expression level. The amount of transfected expression plasmid for HiBiT-tagged fVAMP2 varied (5 to 50 ng) while the amounts of transfected expression plasmids for the t-SNAREs were kept constant. The v-cells expressing HiBiT-tagged-fVAMP2 were mixed with t-cells and cell-cell fusion efficiency was evaluated by measuring the activity of reporter Fluc, followed by HiBiT detection assay to measure the expression level of HiBiT-tagged fVAMP2. (**D**) Inhibition of cell-cell fusion by *cis*-expression of fVAMP2 in t-cells.

**Fig. S2.**
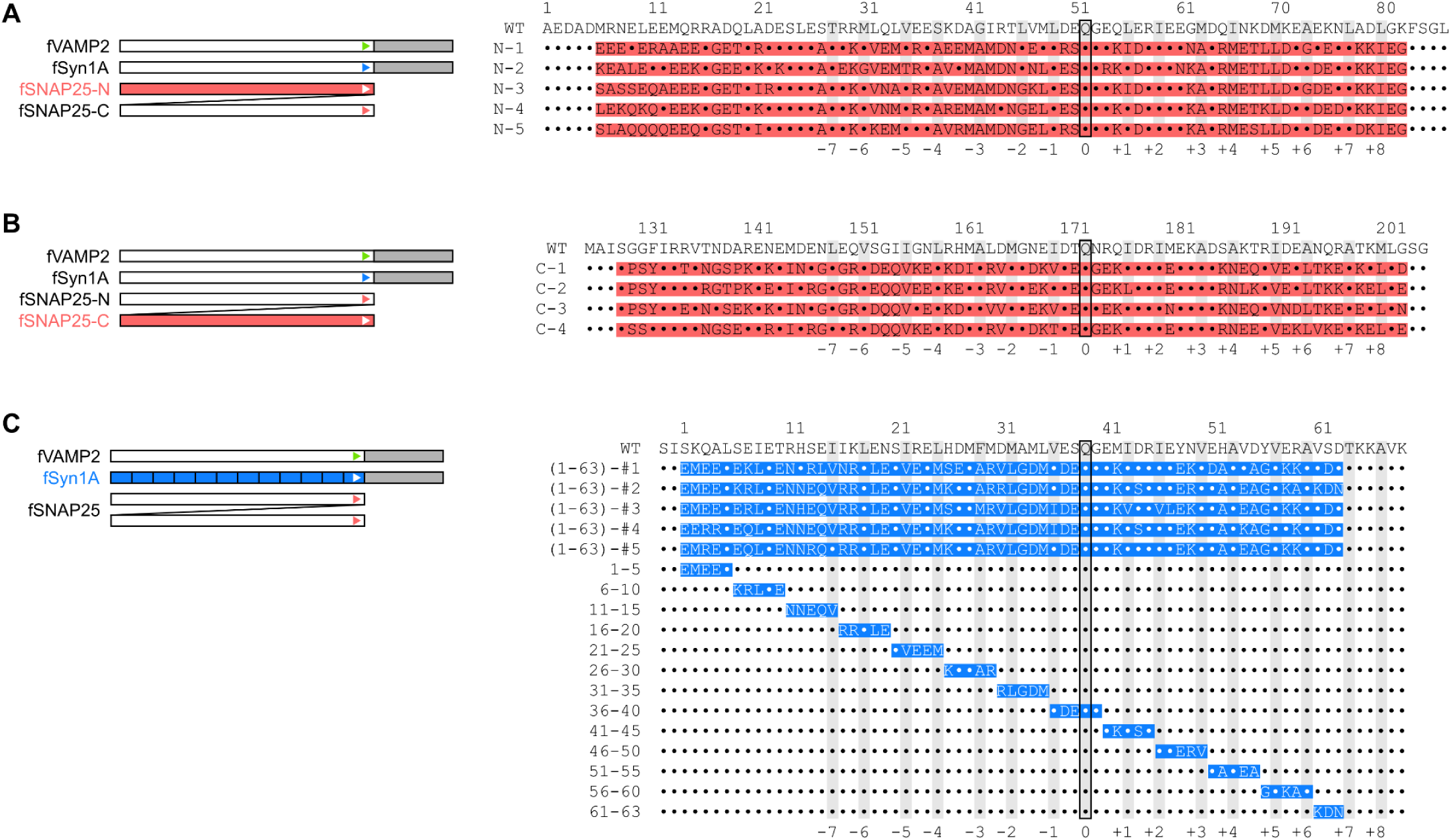
Sequence alignment of redesigned SNAP25 and Syn1A. (**A**) Sequence alignment of the N-terminal coiled-coil domains of WT and redesigned SNAP25 proteins. (**B**) Sequence alignment of SNAP25 with redesigned C-terminal coiled coil domain. (**C**) Sequence alignment of redesigned Syn1A. In all panels, redesigned segments are highlighted in color. The identities of redesigned residues are provided, while WT residues are represented as dots. Gray columns indicate positions at the inner core of each layer of the coiled-coil, and the ionic layer is outlined in black.

**Fig. S3.**
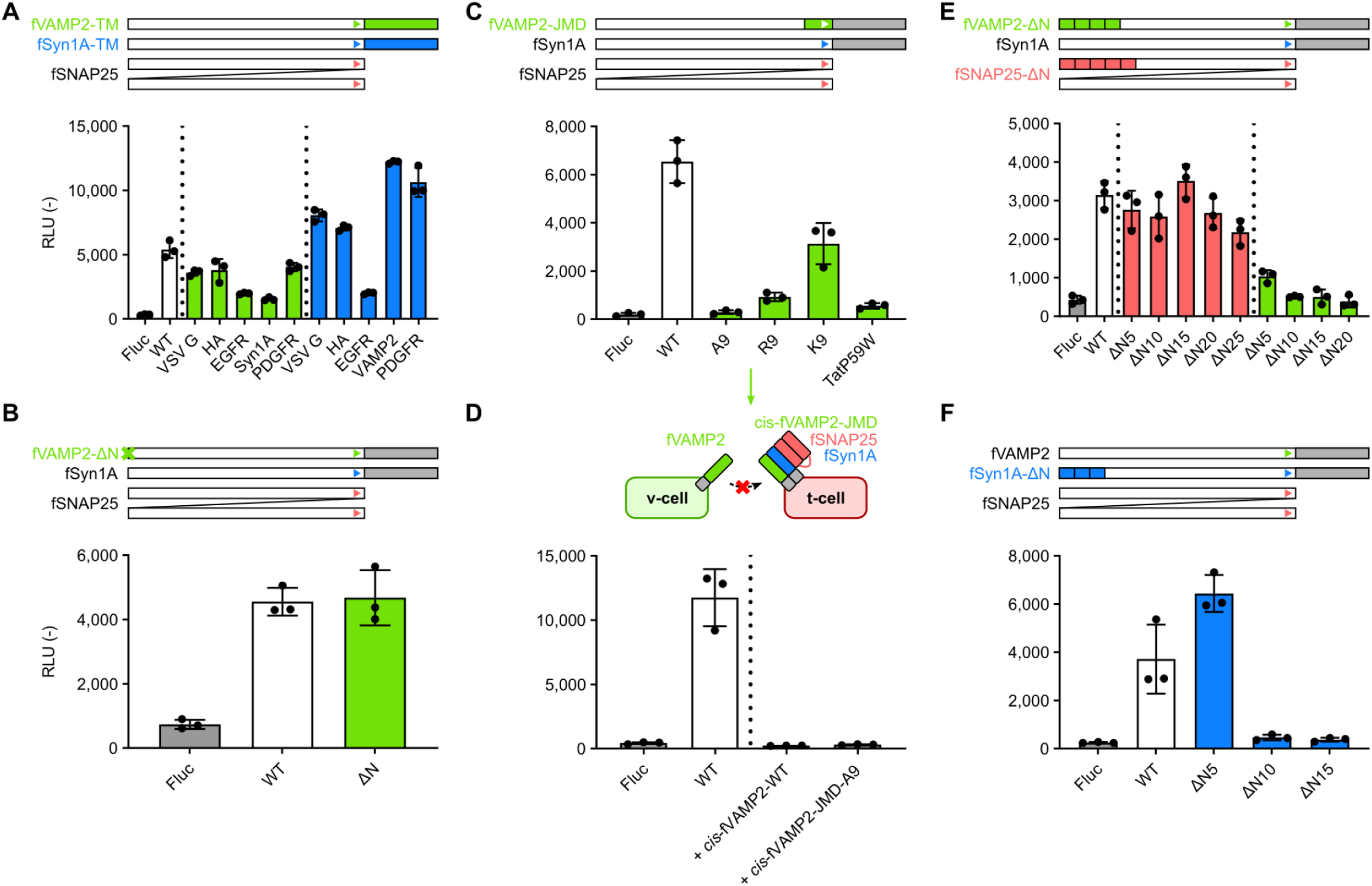
Fusion activity of redesigned three-component fusogens. (**A**) Effects on substitution of the transmembrane domain (TMD) of fVAMP2 and fSyn1A. The native TMD was replaced with non-native TMDs which are derived from vesicular stomatitis virus G protein (VSV-G), influenza virus hemagglutinin (HA), epidermal growth factor receptor (EGFR), or platelet-derived growth factor receptor (PDGFR). Also, TMDs of fVAMP2 and fSyn1A were swapped and tested in a cell-cell fusion assay. (**B**) Truncation of N-terminal Pro-rich unstructured domain of fVAMP2. (**C**) Mutations at the juxtamembrane domain (JMD) of fVAMP2. Mutant sequences are; A9, KLAAAAAAAAAK; R9, KLRRRRRRRRRK; K9, KLKKKKKKKKKK; TatP59, KLGRKKRRQRRRPWQK. (**D**) Blockade of cell-cell fusion by *cis*-expression of fVAMP2-WT and fVAMP2-JMD-A9 mutant in t-cells. (**E**) Truncation of N-terminal region of fVAMP2 and fSNAP25. Five to twenty-five amino acids were truncated from the N terminus of fVAMP2 or fSNAP25 and evaluated in a cell-cell fusion assay. (**F**) Truncation of N-terminal region of fSyn1A. Five to fifteen amino acids were truncated from the N terminus of fSyn1A and evaluated in a cell-cell fusion assay.

**Fig. S4.**
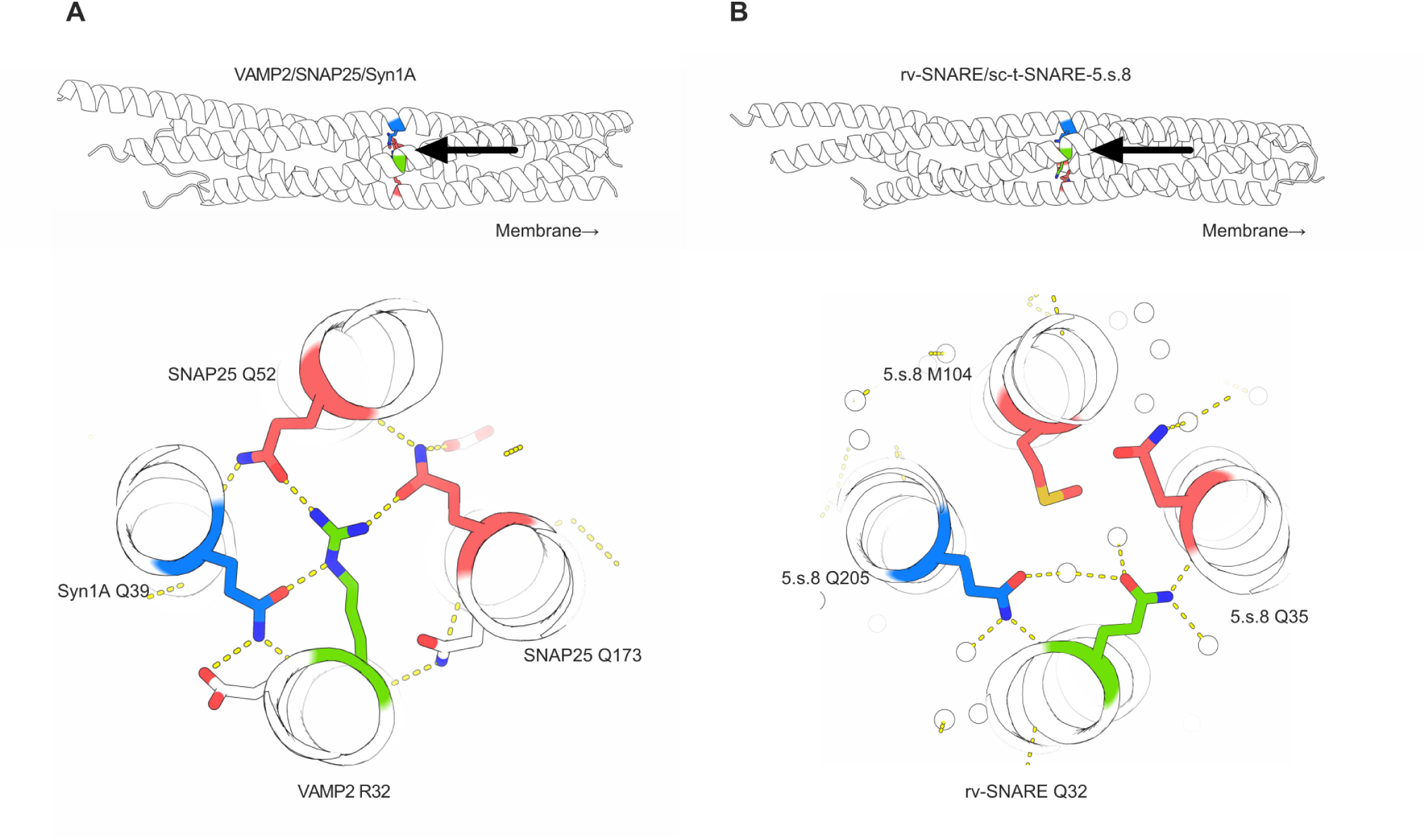
Crystal structure of the two-component fusogen. (**A**) *Top,* the ionic layer in the crystal structure of the hnSNARE complex (PDB ID: 1sfc) is indicated in color (as in **Fig. 1A**). (*Bottom*) The ionic layer is viewed from the membrane side, as indicated by the large arrows above. Yellow dashed lines represent hydrogen bonds. (**B**) The same layer in the crystal structure of a synthetic two-component fusogen (rv-SNARE and sc-t-SNARE-5.s.8). Water molecules are represented as white spheres.

**Fig. S5.**
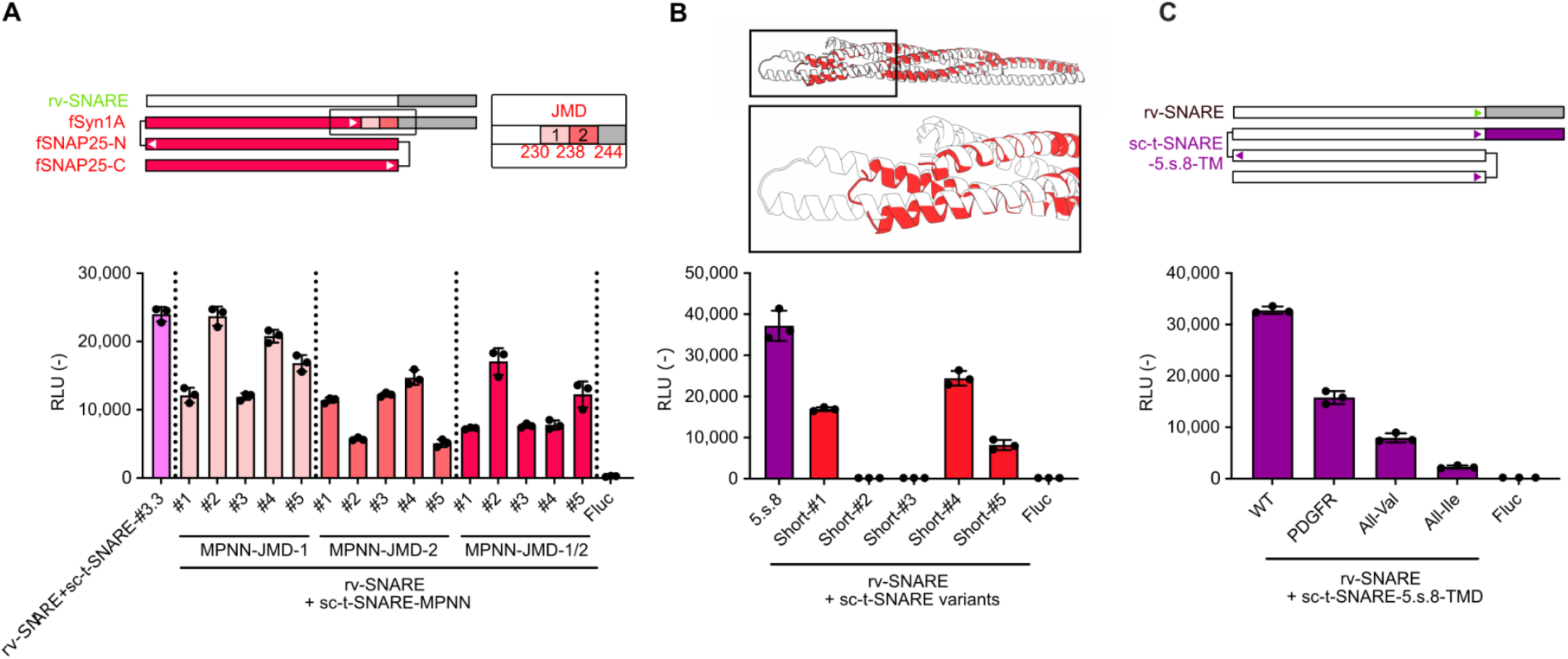
Variants of sc-t-SNAREs. (**A**) Fusion efficiency of sc-t-SNARE variants with redesigned coiled coil domains and JMD. The amino acid sequence of these new designs were redesigned by ProteinMPNN from parental sc-t-SNARE-#3.3 including JMD-1 (N-terminal half of JMD) and/or JMD-2 (C-terminal half of JMD). As a control, sc-t-SNARE-#3.3 which has native Syn1A-JMD sequence was used. (**B**) The sc-t-SNARE variants with truncated SNAP25-N portion. Superposition of the predicted structures of rv-SNARE and sc-t-SNARE-5.s.8 (white), and the redesigned shorter variant of sc-t-SNARE (red). The membrane fusion efficiency of five truncated sc-t-SNARE variants (lower). As a control, sc-t-SNARE-5.s.8 was used. (**C**) Variants of sc-t-SNARE-5.s.8 with non-native TMD sequences including PDGFR, poly-Val, and poly-Ile.

**Fig. S6.**
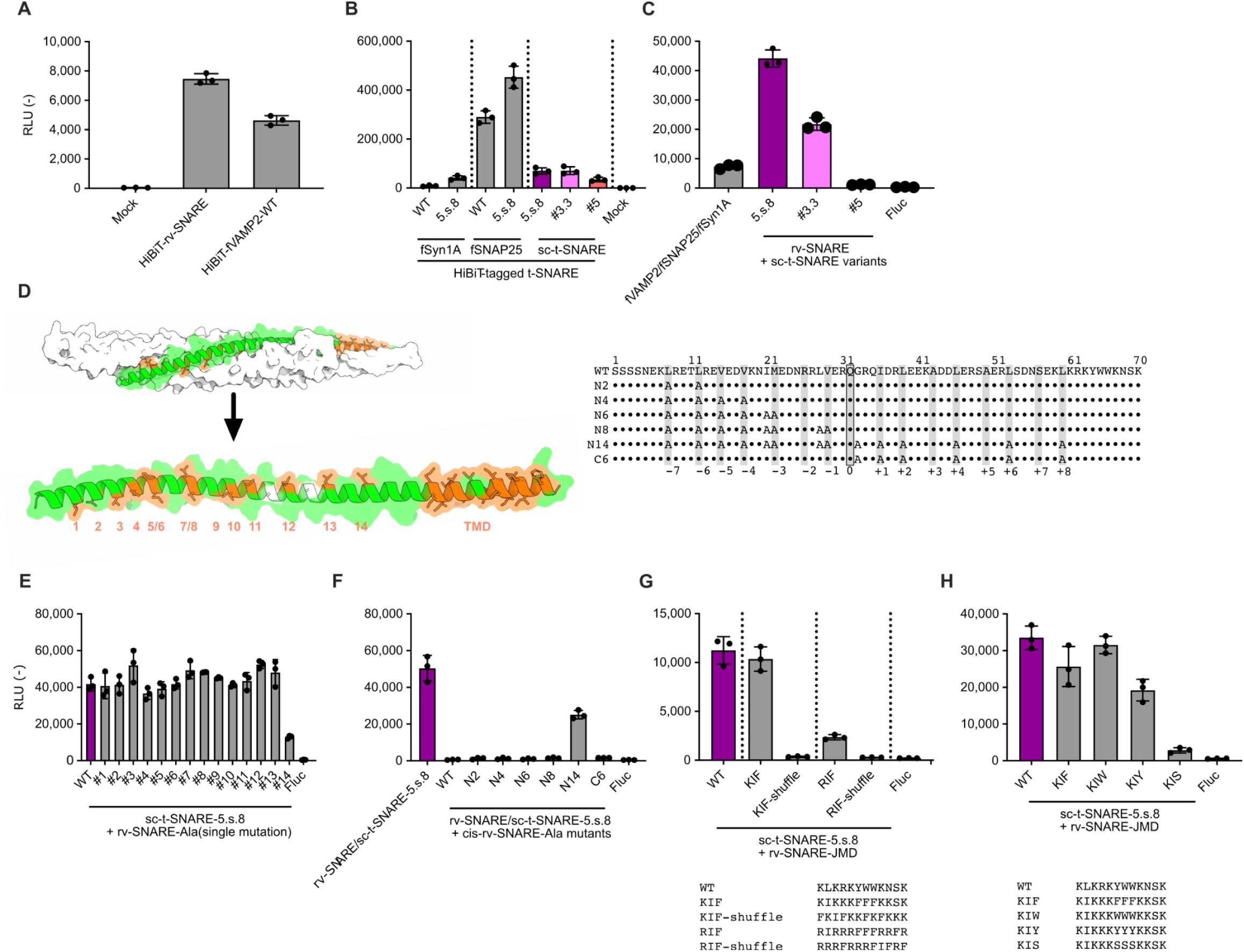
Characterization of two-component fusogens. (**A**) Expression level of HiBiT-tagged fVAMP2 and rv-SNARE. HEK293T cells expressing synthetic fusogens were lysed and mixed with recombinant LgBiT protein and Nluc substrate, followed by measurement of luminescence signal. (**B**) Expression level of HiBiT-tagged native and redesigned t-SNAREs and sc-t-SNARE. The amino acid sequence of redesigned t-SNAREs are derived from sc-t-SNARE-5.s.8. (**C**) Membrane fusion activity of sc-t-SNARE variants. (**D**) Positions of hydrophobic residues of rv-SNARE at the interface to the sc-t-SNARE. Predicted complex structure (left) of rv-SNARE (green) and sc-t-SNARE5.s.8 (white). Hydrophobic residues of rv-SNARE are highlighted in orange and numbered, and alanine residues at the interface are highlighted in white. Sequence alignment of rv-SNARE variants with Ala mutations at the hydrophobic residues (right). Layers are highlighted as shading and numbered from -7 to +8. (**E**) Effect of single Ala mutation in rv-SNARE on cell-cell fusion activity. (**F**) Effect of multiple Ala mutations in rv-SNARE on cell-cell fusion activity when expressed *in cis* to sc-t-SNARE-5.s.8. Interface hydrophobic residues are mutated to Ala; a total of 2, 4, 6, 8, and 14 alanines from the N-terminus or 6 alanines from the C-terminus. (**G**) Effect of JMD sequence in rv-SNARE on cell-cell fusion activity. Note that WT-JMD sequence of rv-SNARE has 1 mutation at the 11th position (native JMD sequence of VAMP2 is KLKRKYWWKN<u>L</u>K). (**H**) Variants of rv-SNARE with synthetic JMD sequence.

**Fig. S7.**
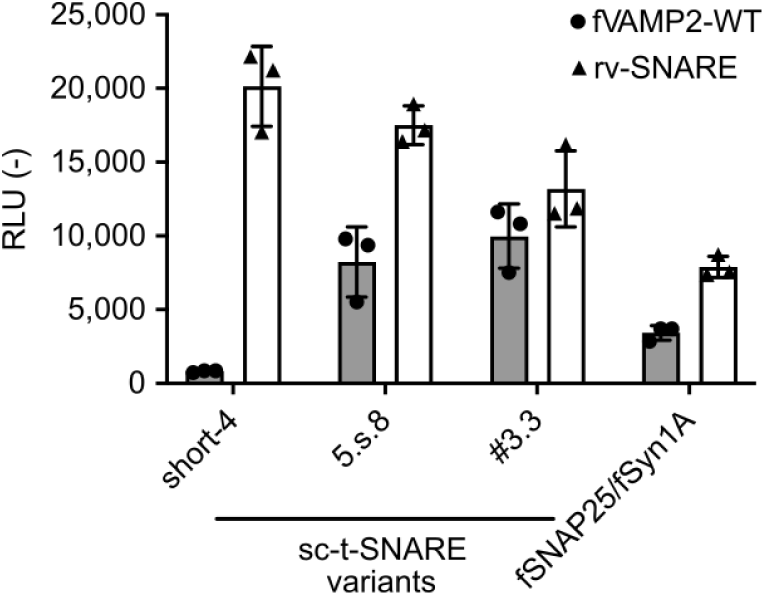
Compatibility of v-SNARE and sc-t-SNARE variants. Cell-cell fusion activity of various combinations of synthetic fusogens. The v-SNARE variants include fVAMP2-WT and rv-SNARE, and the t-SNARE variants include sc-t-SNARE-#3.3, sc-t-SNARE-5.s.8, and sc-t-SNARE-short-4.

**Fig. S8.**
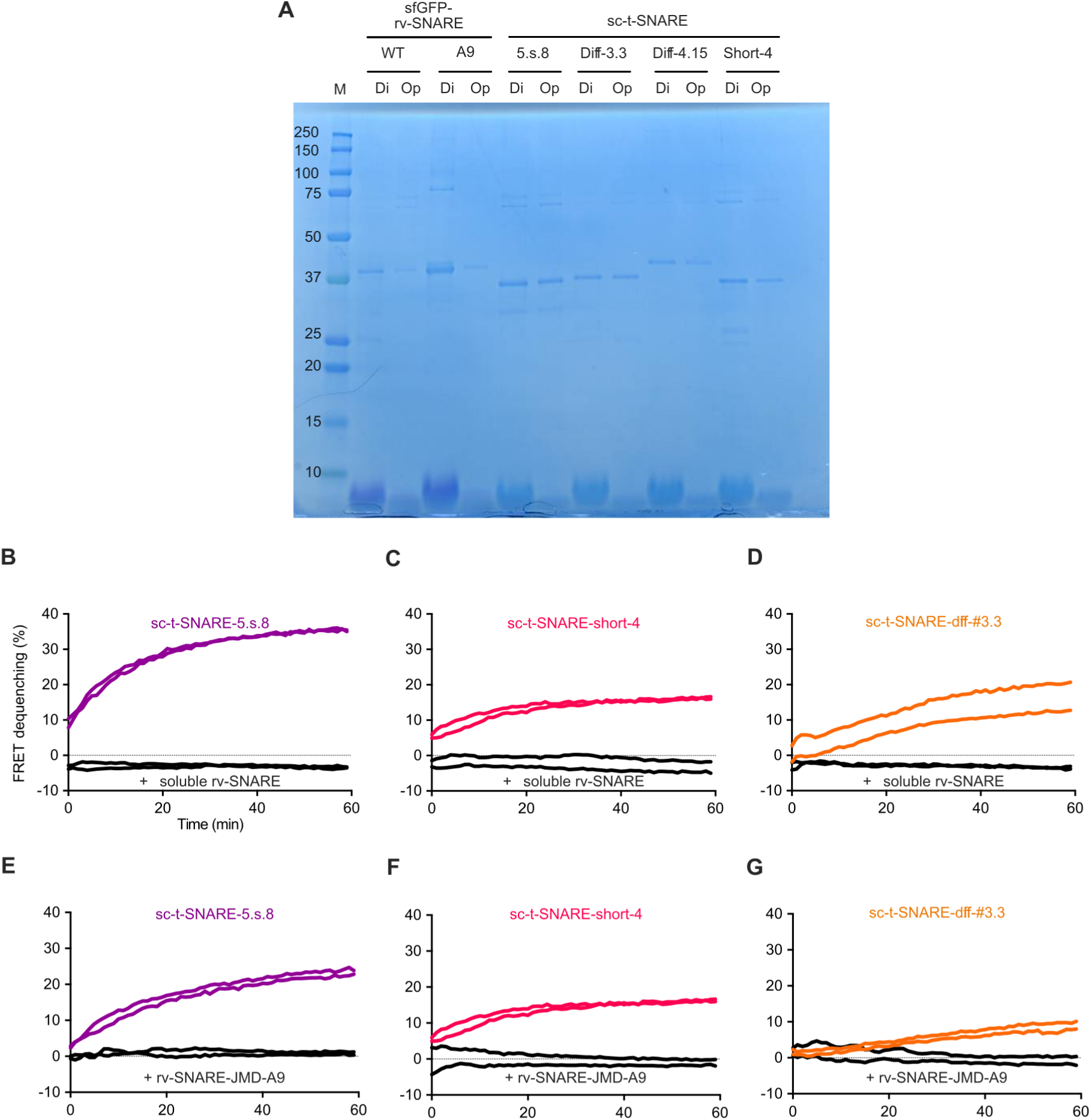
Reconstitution of synthetic fusogens into liposomes. (**A**) Proteoliposomes containing rv-SNARE (WT or JMD-A9 mutant) or sc-t-SNARE variants were analyzed by SDS–PAGE and visualized by Coomassie staining. “Di” and “Op” indicate proteoliposome samples collected after dialysis and OptiPrep isolation, respectively. (**B**-**D**) Inhibition of liposomal fusion by excess soluble rv-SNARE. Liposomes reconstituted with sc-t-SNARE variants were mixed with liposomes containing rv-SNARE in the presence (black) or absence (colored) of soluble rv-SNARE. FRET signal was monitored for 60 min at 37°C. (**E**-**G**) Effect of the rv-SNARE JMD mutation on fusion activity. Liposomes with sc-t-SNARE variants were fused with rv-SNARE liposomes carrying either WT-JMD (colored) or A9-JMD (black).

**Fig. S9.**
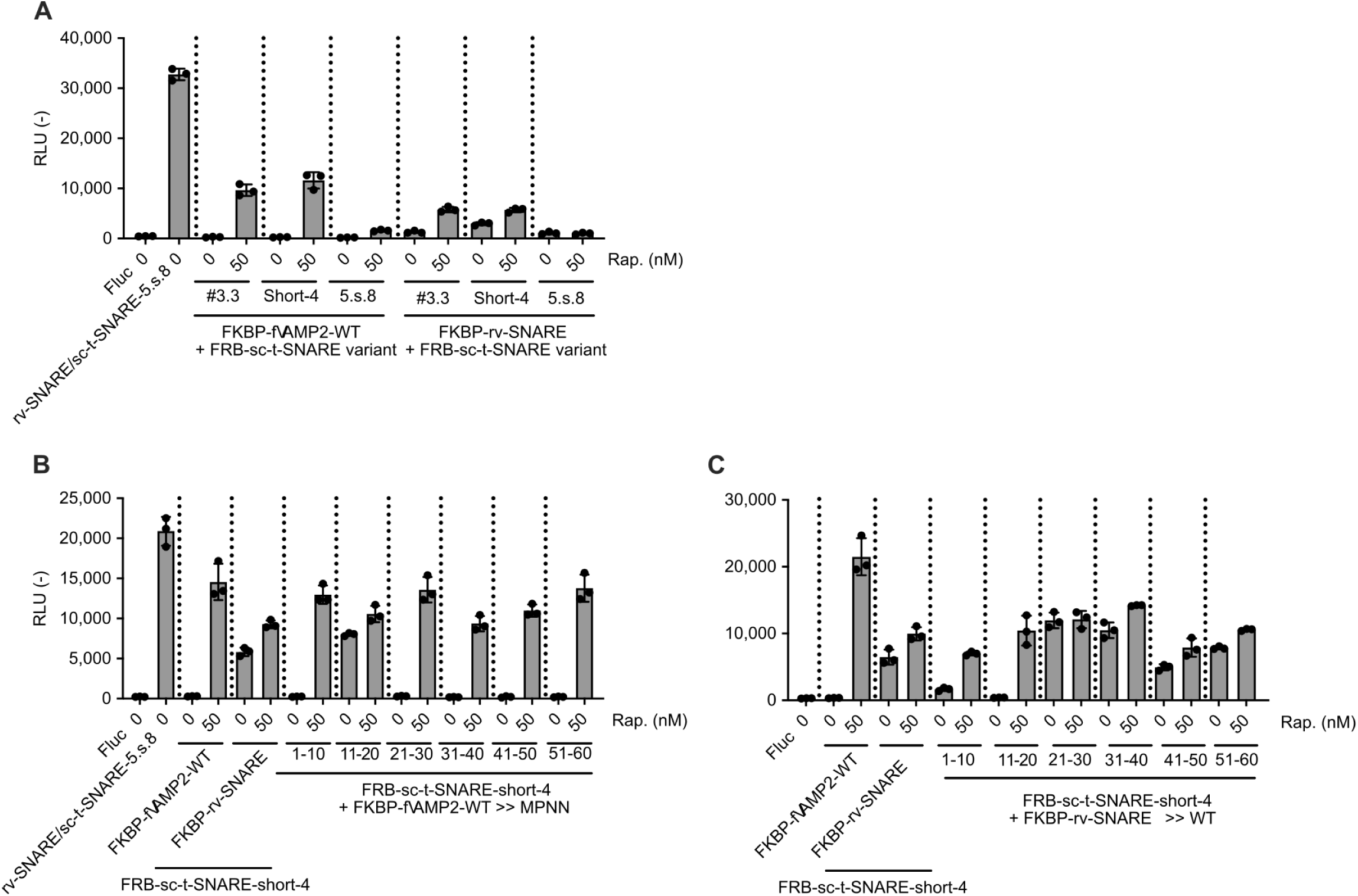
Optimization of inducible two-component fusogens. (**A**) Comparison of FKBP-fused v-SNARE variants and FRB-fused sc-t-SNARE variants on inducible cell-cell fusion activity. The v-cell and t-cells were mixed and incubated in the presence of rapamycin (0 or 50 nM). (**B**) Swapping of the native VAMP2 sequence of FKBP-fused v-SNARE to MPNN-redesigned sequences. (**C**) Swapping of the redesigned v-SNARE sequence of FKBP-fused v-SNARE to VAMP2-native sequences.

**Fig. S10.**
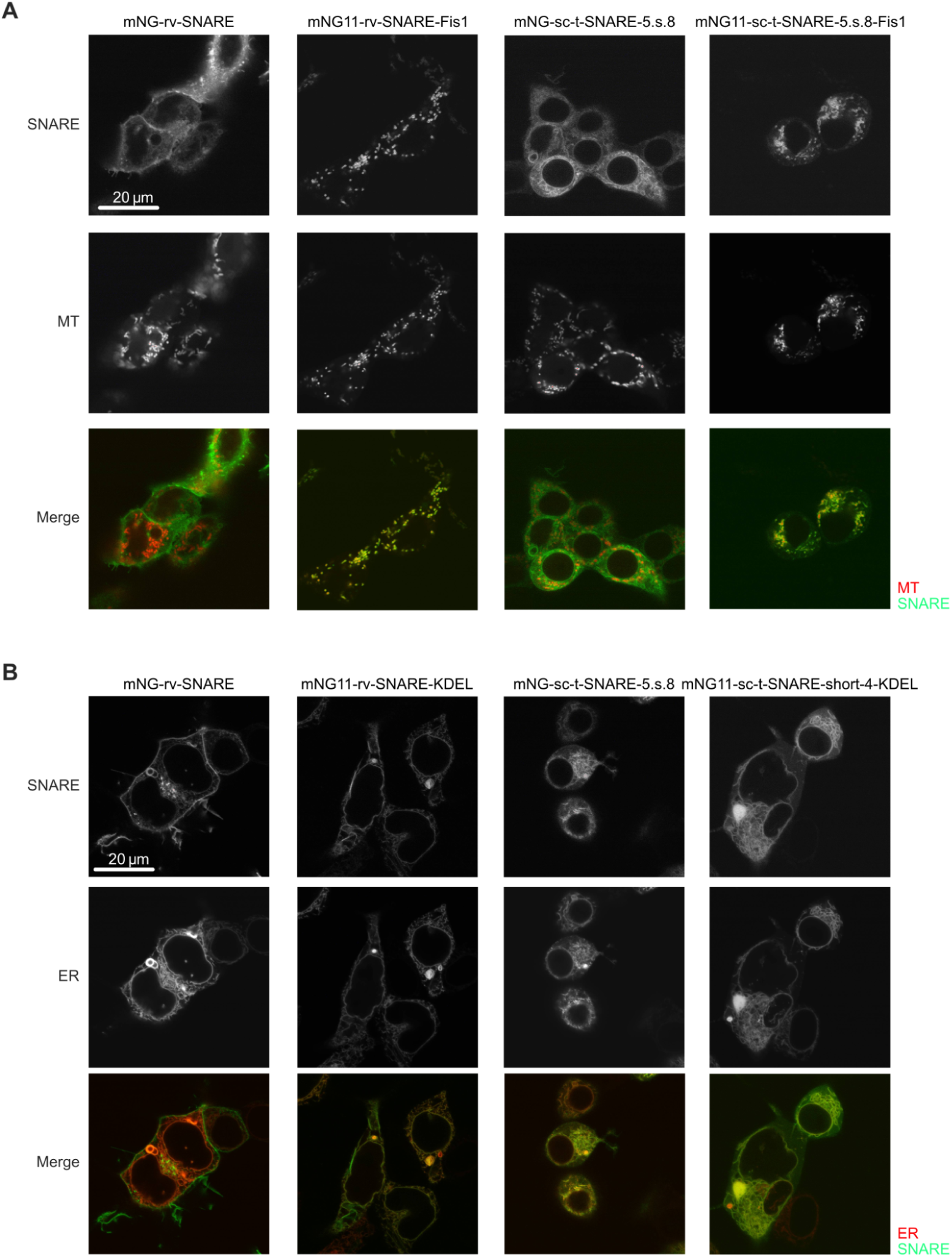
Expression of synthetic fusogens targeting specific organelles in HEK293T cells. (**A**) Confocal images of HEK293T cells expressing synthetic fusogens with the C-terminal mitochondrial outer membrane (MOM)-targeting signal derived from Fis1. Synthetic fusogens were fused with either mNeonGreen or split mNeonGreen tag (mNG2_11_) at the N terminus, and visualized with or without the large subunit of split mNG (mNG2_1-10_). Mitochondria (MT) were labeled with MT matrix-localized mScarlet (4×mts-mScarlet-I) (*3*). (**B**) Localization of ER-targeting synthetic fusogens in HEK293T cells. Synthetic fusogens were fused with mNG tag at the N terminus as in panel (A), and also included a KDEL sequence at the C terminus for ER retention. ER was labeled with KDEL-fused mScarlet-I (ER-mScarlet-I).

**Fig. S11.**
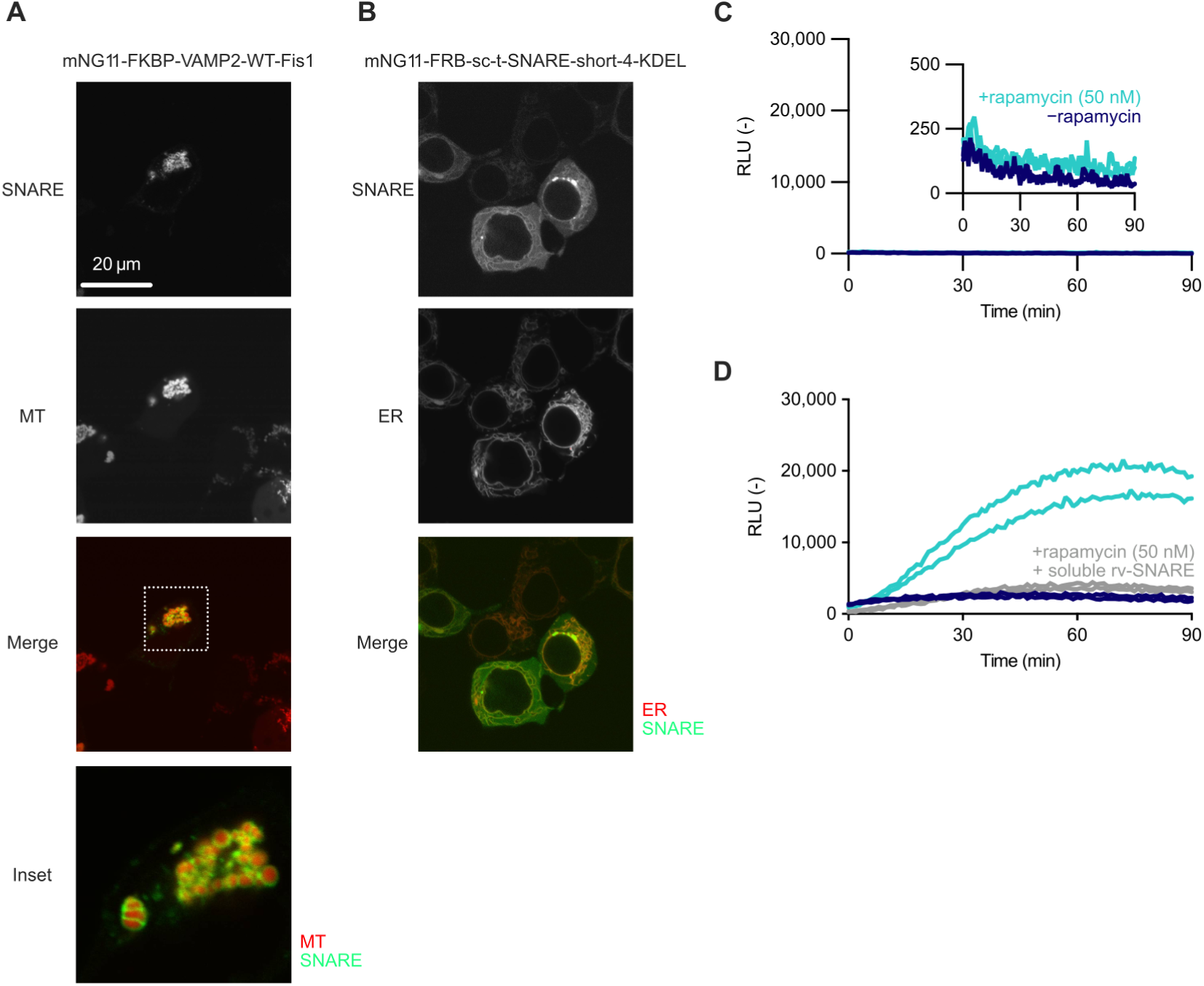
Fusion of endoplasmic reticulum and mitochondrial outer membrane by chemically inducible fusogens. (**A**) Confocal images of HEK293T cells expressing mNG11-FKBP-VAMP2-WT-Fis1 and MT matrix marker. Magnified image indicates ring-like shape, suggesting the localization of mNG11-FKBP-VAMP2-WT-Fis1 on MOM. (**B**) Localization of mNG11-FRB-sc-t-SNARE-short-4-KDEL and ER marker. (**C**) and (**D**) Rapamycin-dependent fusion of ER and MT by synthetic fusogen measured by split Nluc. (**C**) ER-MT fusion with the HiBiT probe localized in the mitochondrial matrix and the LgBiT probe localized in ER lumen. (**D**) The inhibition of fusion by soluble rv-SNARE that binds to sc-t-SNARE and blocks fusion.

**Table S1.** Plasmids, reagents, and other materials.

|  | Name | Manufacturer/source | Identifier |
| --- | --- | --- | --- |
| Plasmids | pCMV vector for golden gate cloning | This study | V338 |
|  | pcDNA3.1 vector for golden gate cloning | This study | V337 |
|  | pET29b vector for golden gate cloning | This study | V366 |
|  | pCMV-Fluc | This study | P20 |
|  | pCMV-T7polymerase | This study | V4 |
|  | pCMV-mNeonGreen | This study | V5 |
|  | pCMV-DsRedEx | This study | V6 |
|  | T7-IRES-NanoLuc | This study | V8 |
|  | T7-IRES-Fluc | This study | V83 |
|  | pCAG-EGFP(HiBiT) | Addgene | #162589 |
|  | pcDNA3.1-FRB-LgBiT | Addgene | #182650 |
|  | pcDNA3.1-hSyncytin-1 | Addgene | #182440 |
|  | 4×mts-mScarlet-I | Addgene | #98818 |
|  | ER-mScarletI | Addgene | #137805 |
| Kits | Nano-Glo® Luciferase Assay System | Promega | N1110 |
|  | Nano-Glo® HiBiT Lytic Detection System | Promega | N3030 |
|  | Nano-Glo® Live Cell Assay System | Promega | N2011 |
|  | Nano-Glo® Dual-Luciferase® Reporter Assay System | Promega | N1610 |
|  | Pur-A-Lyzer(TM) Maxi Dialysis Kit, Maxi 12000 | Sigma Aldrich | PURX12050-1KT |
| Reagents | PEI MAX (linear PEI, MW 40,000) | Polysciences | 24765 |
|  | DrkBiT peptide | Synthesized | - |
|  | Rapamycin | FUJIFILM-Wako | 184-02531 |
|  | DOPC | NOF | MC-8181 |
|  | Cholesterol | Sigma Aldrich | C8667 |
|  | DiD | TCI | D6263-1ML |
|  | Dil | FUJIFILM-Wako | 045-33421 |
|  | *n*-octyl-β-D-glucoside | Nacalai Tesque | 25535-82 |
|  | Opti-Prep | Serumwerk Bernburg AG | 1893 |

**Table S2.**
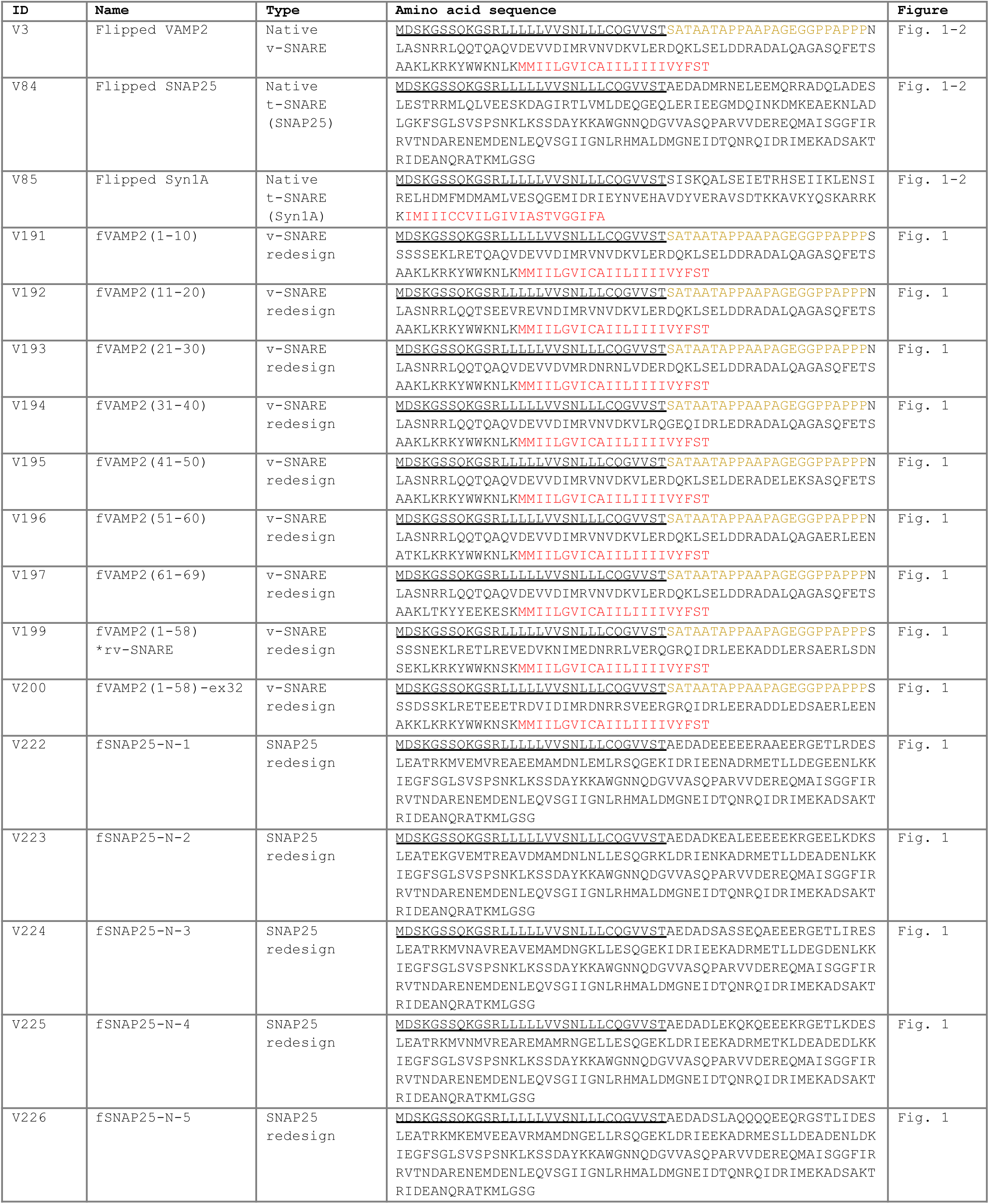
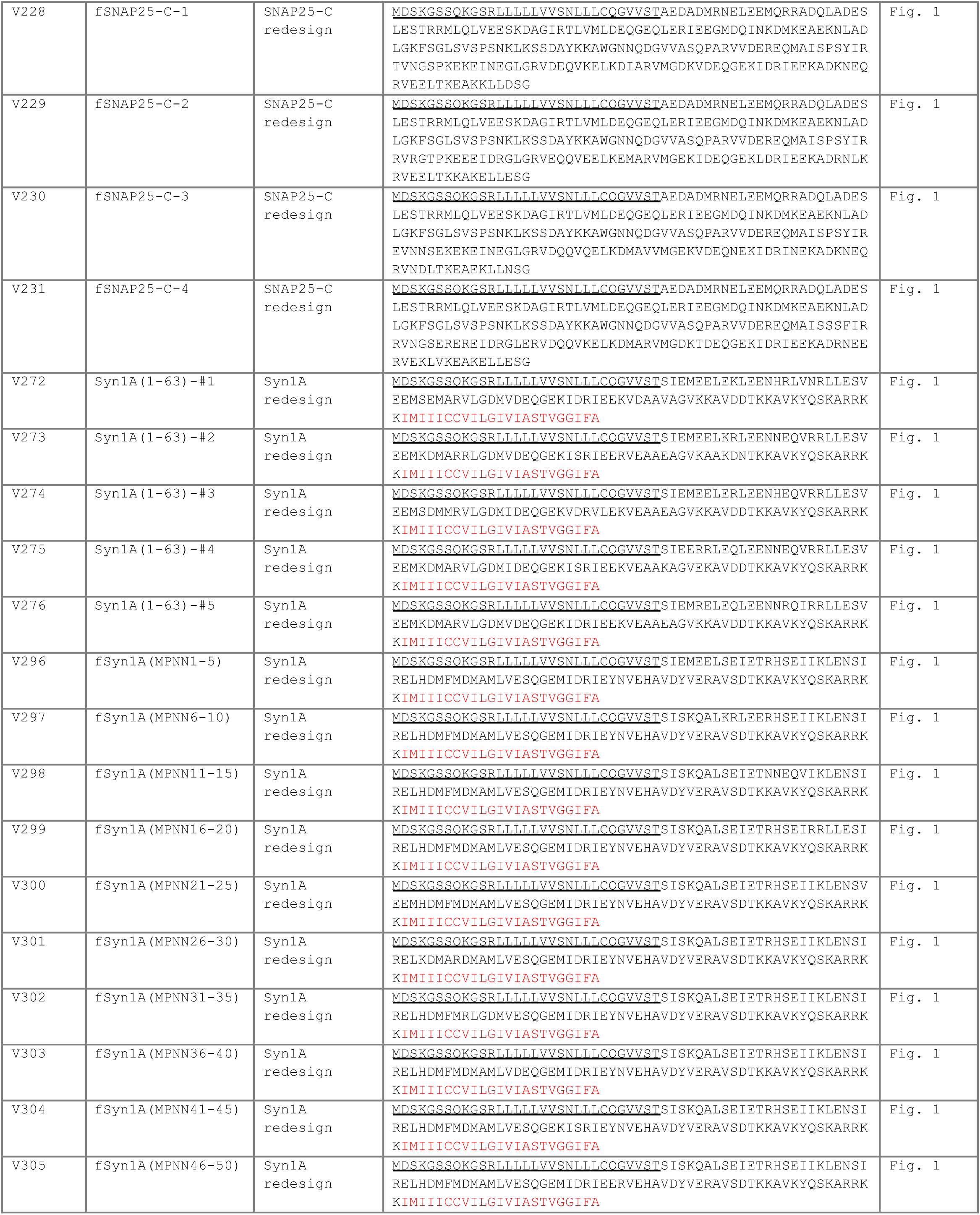
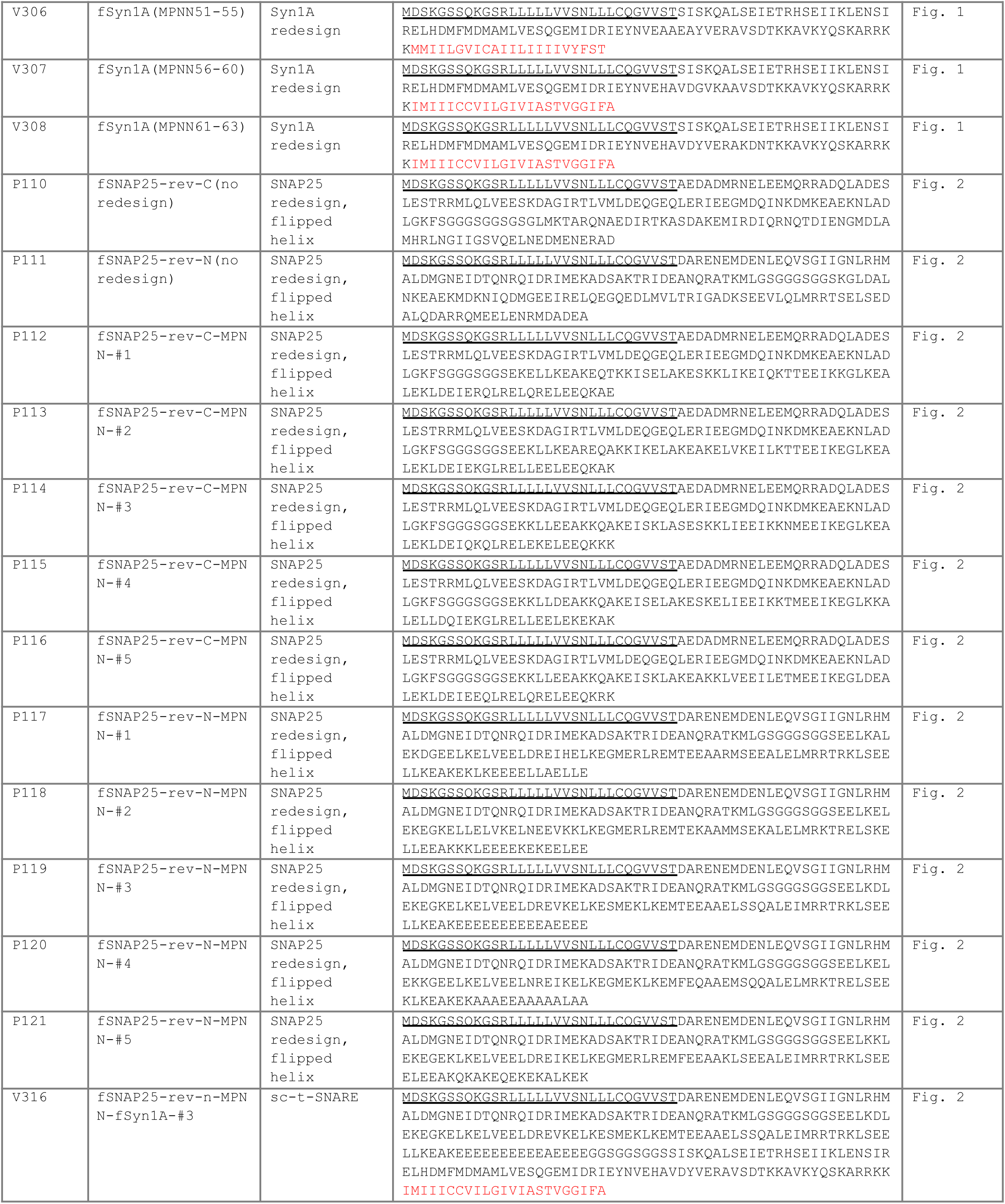
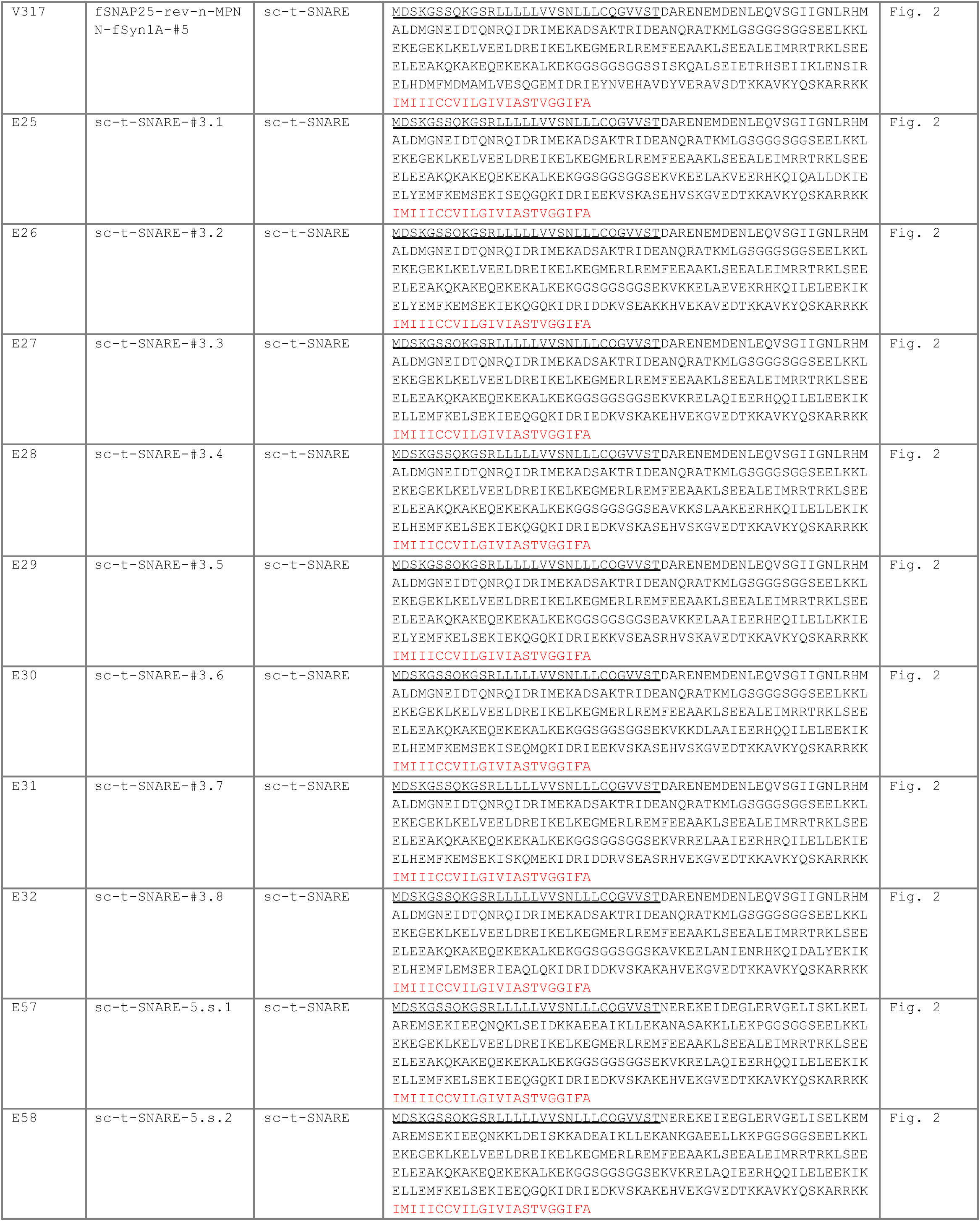
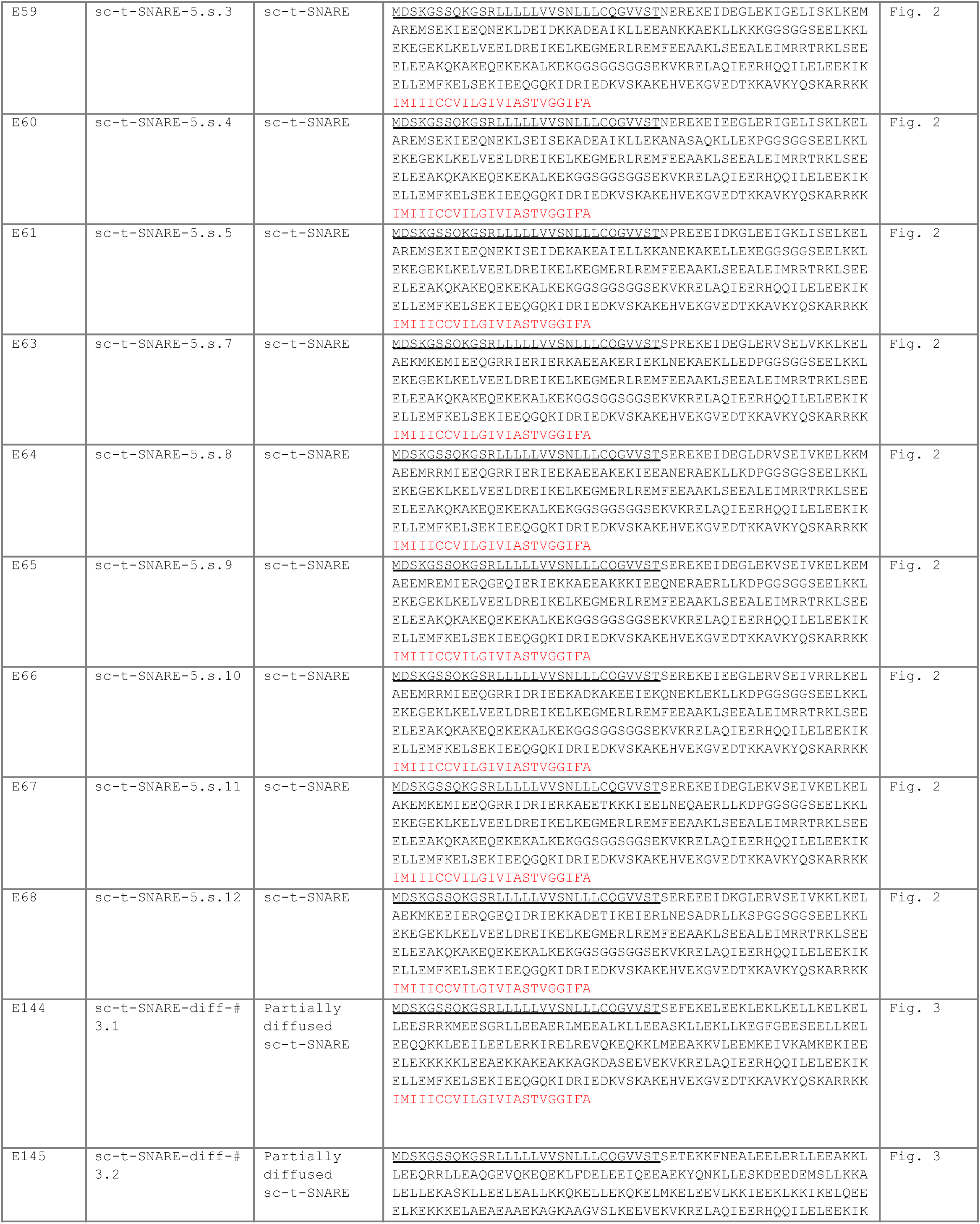
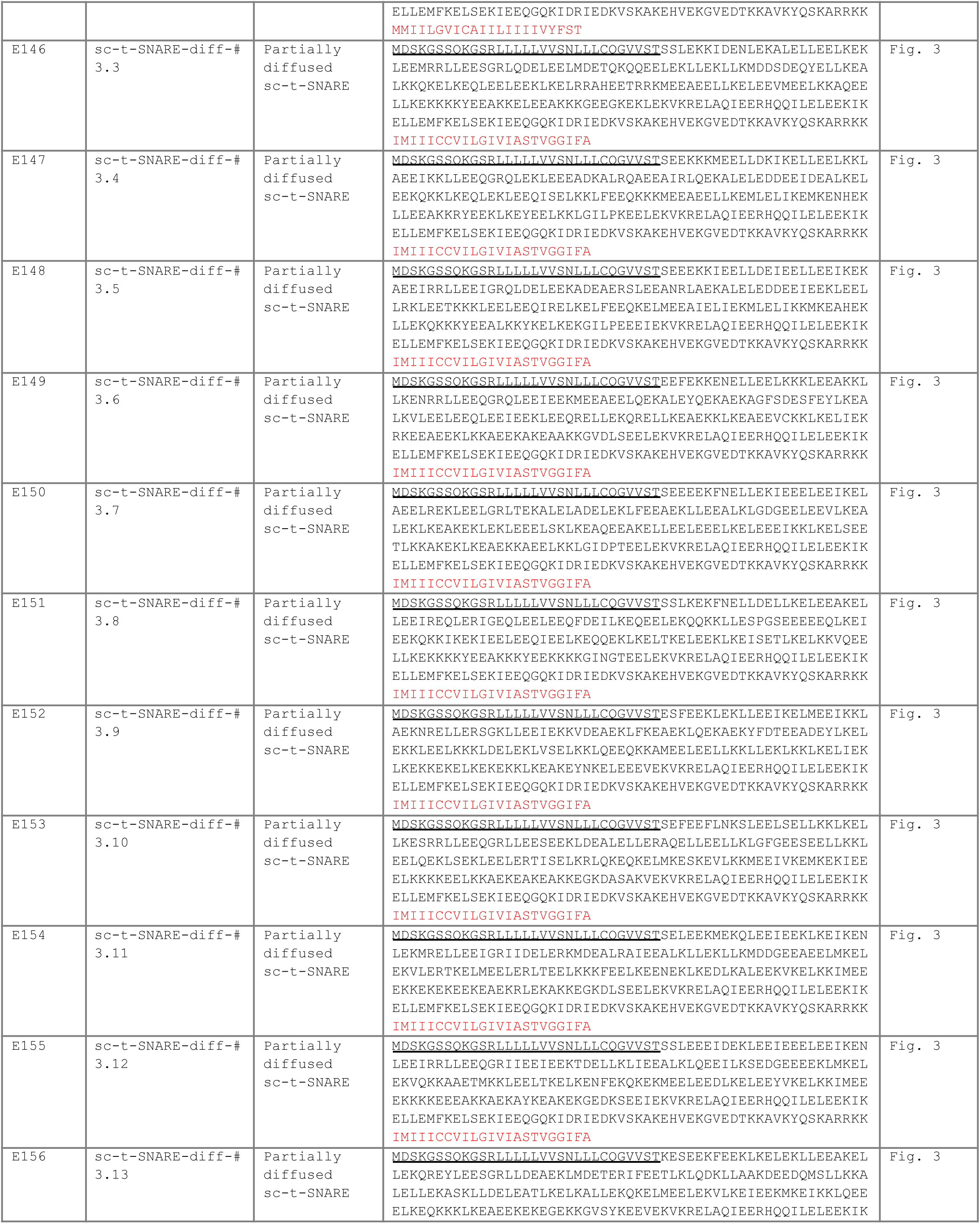
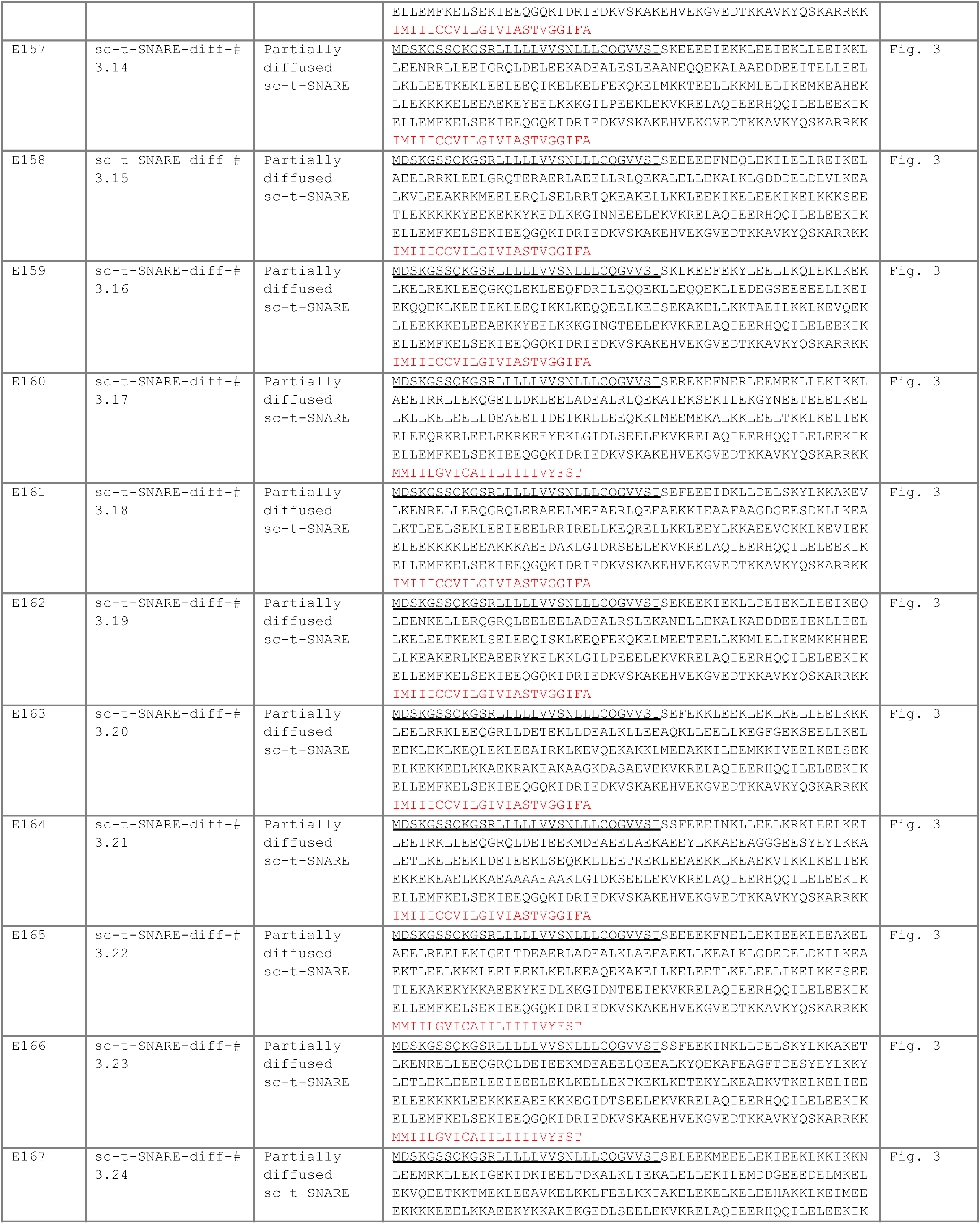
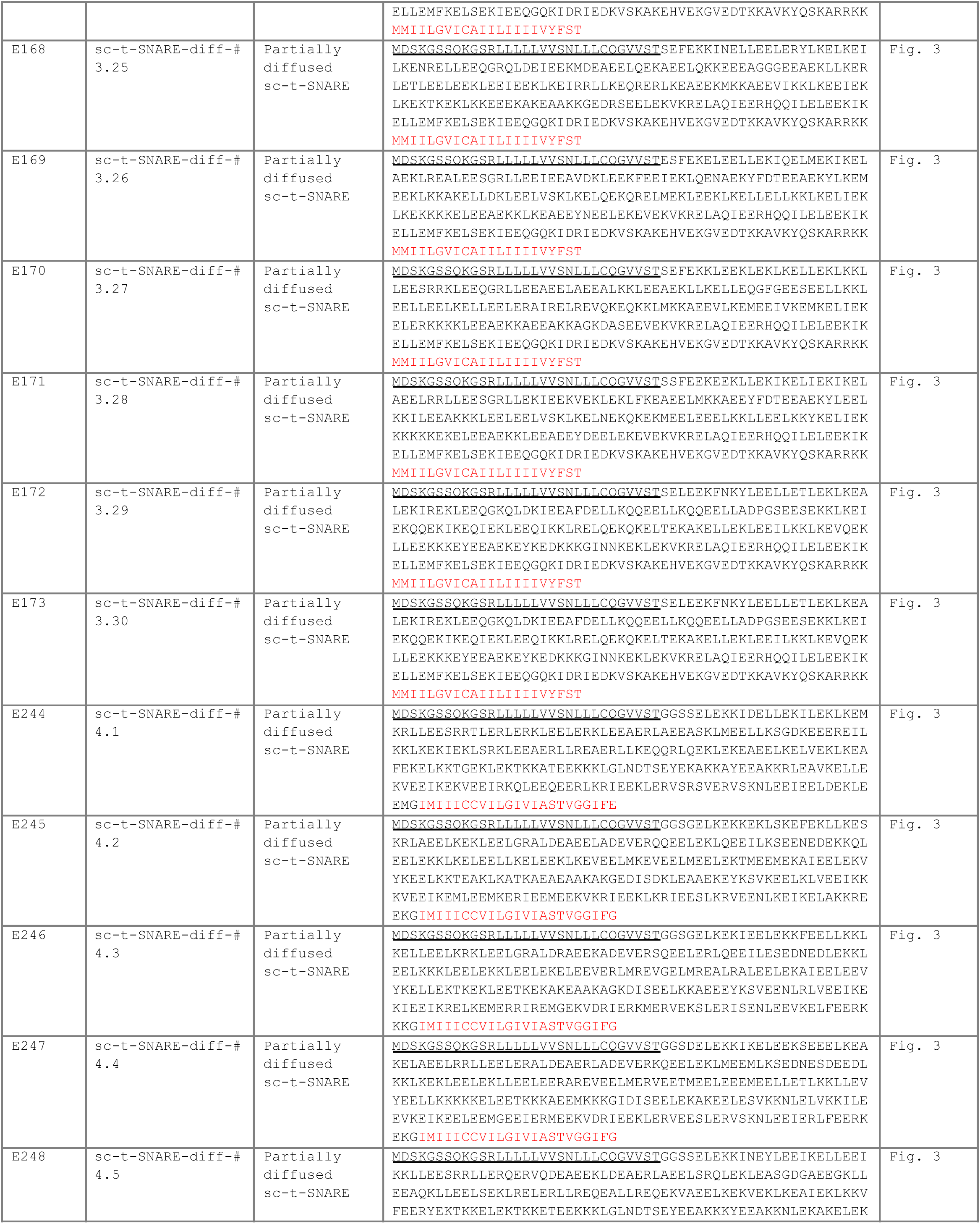
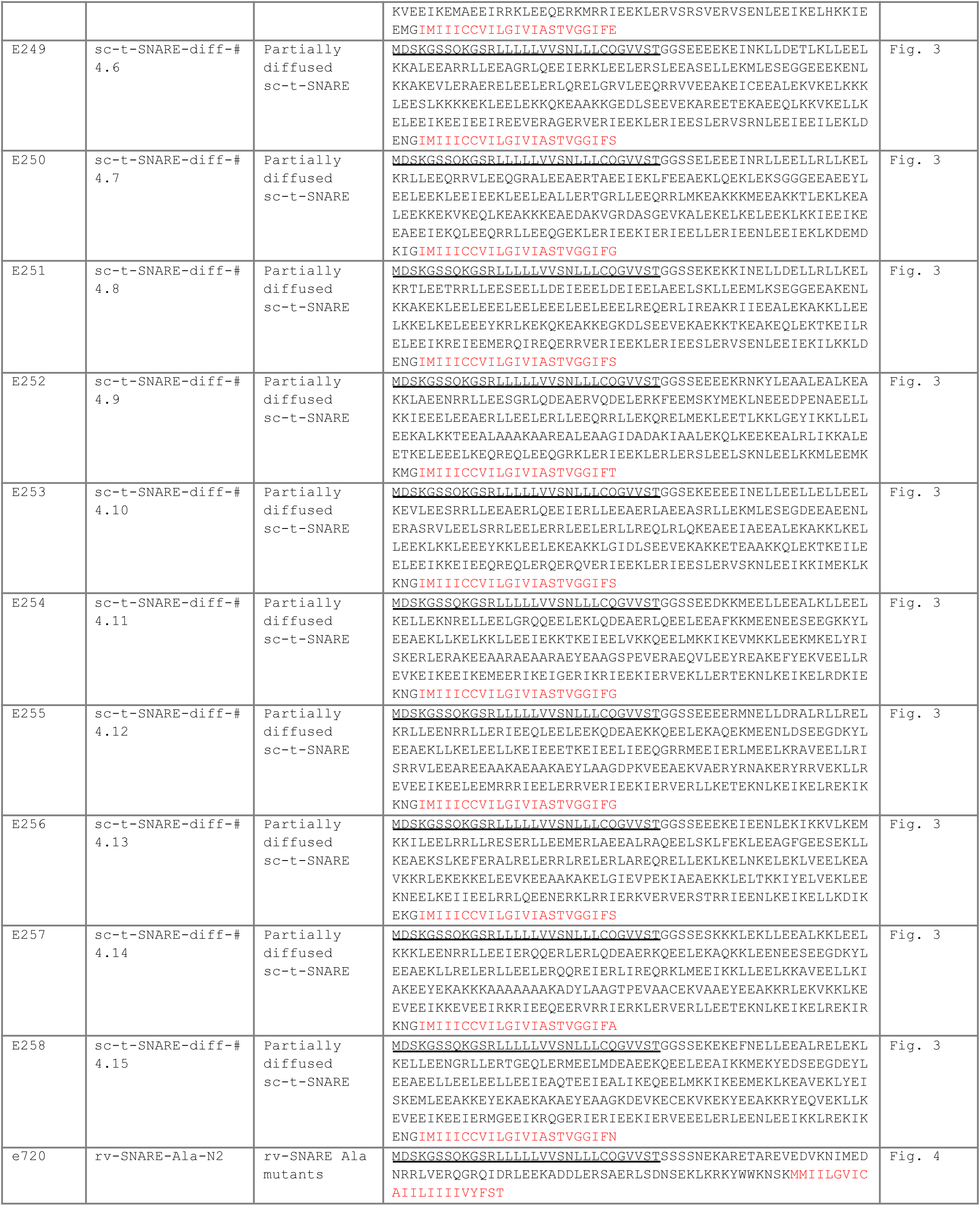
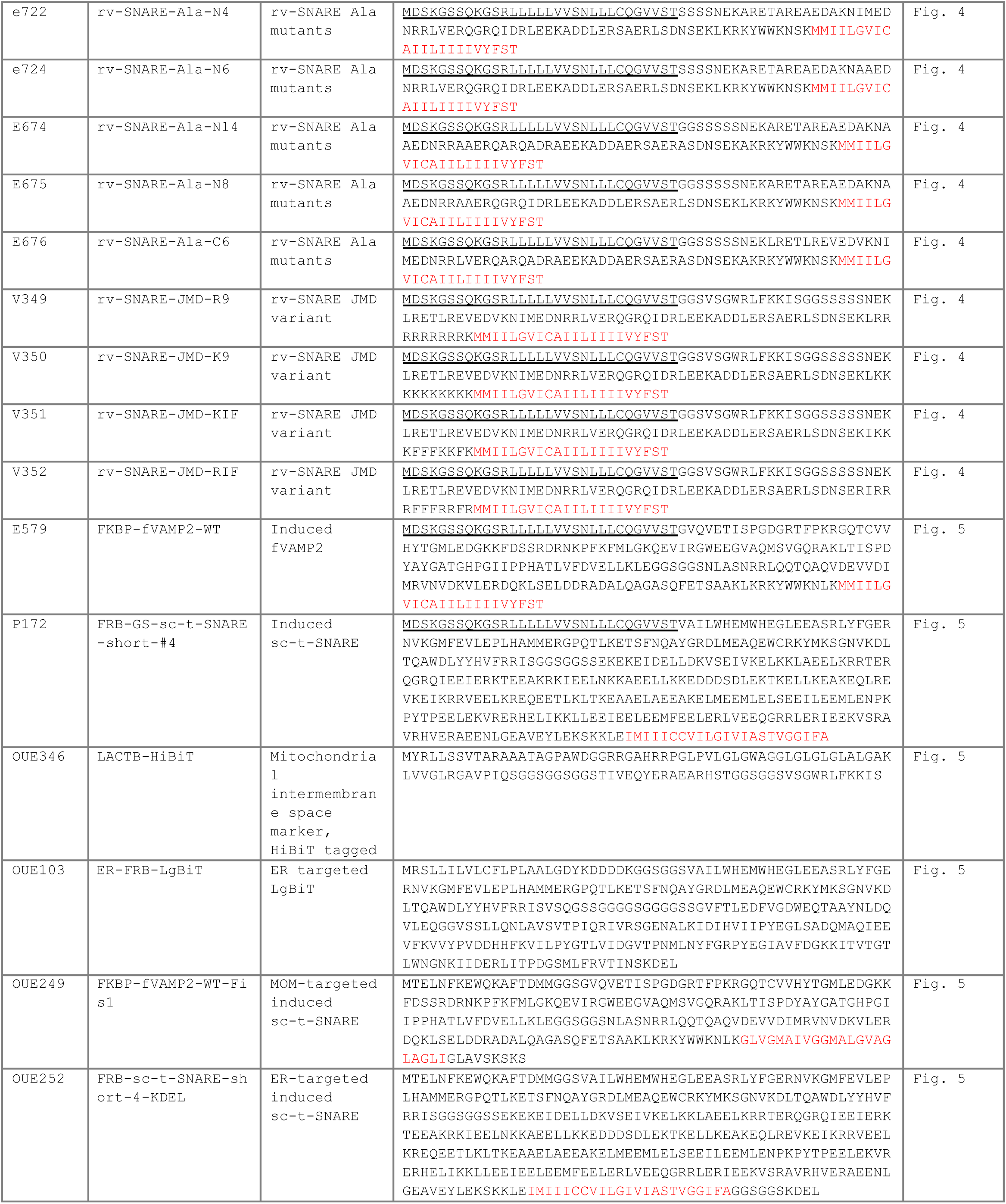
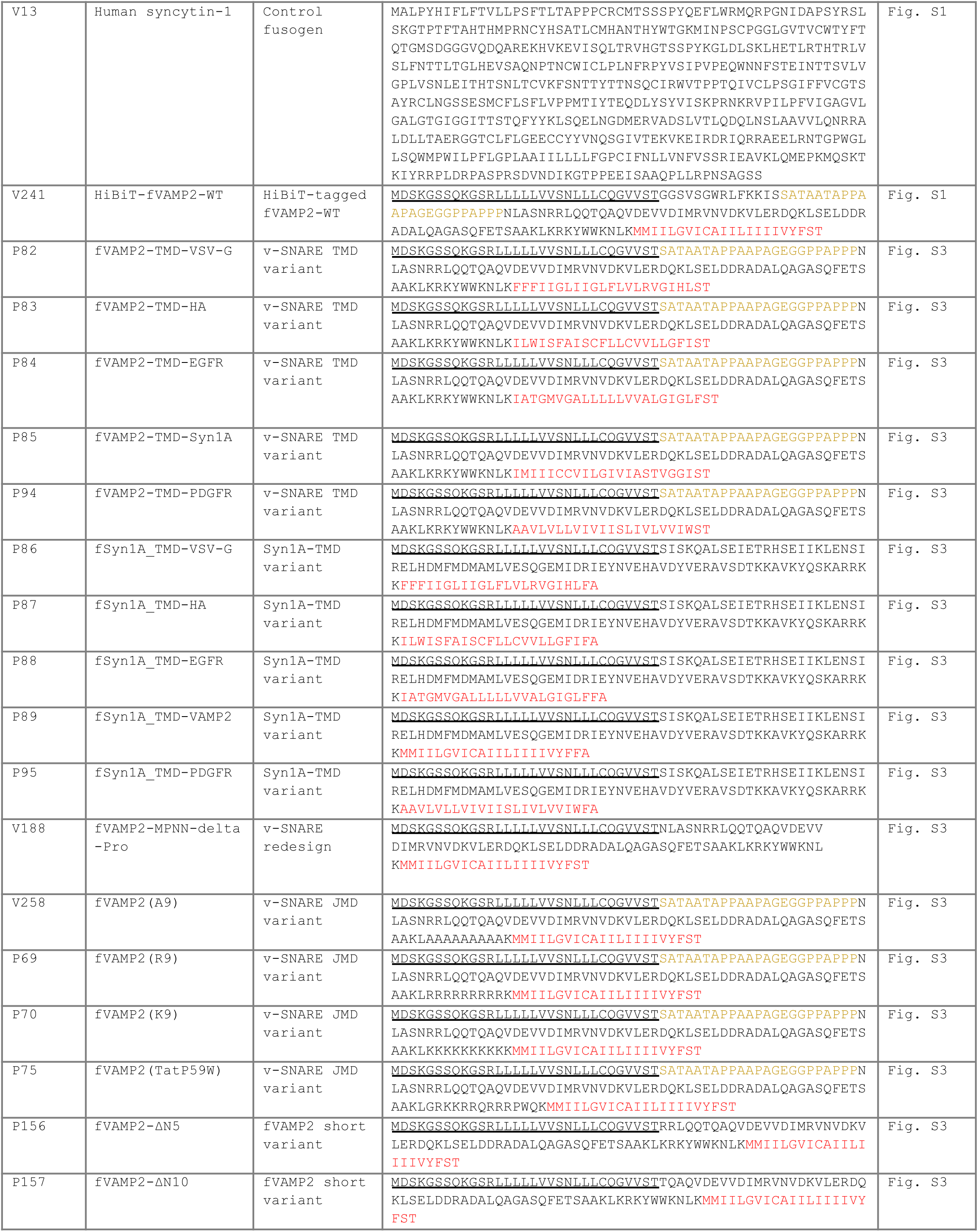
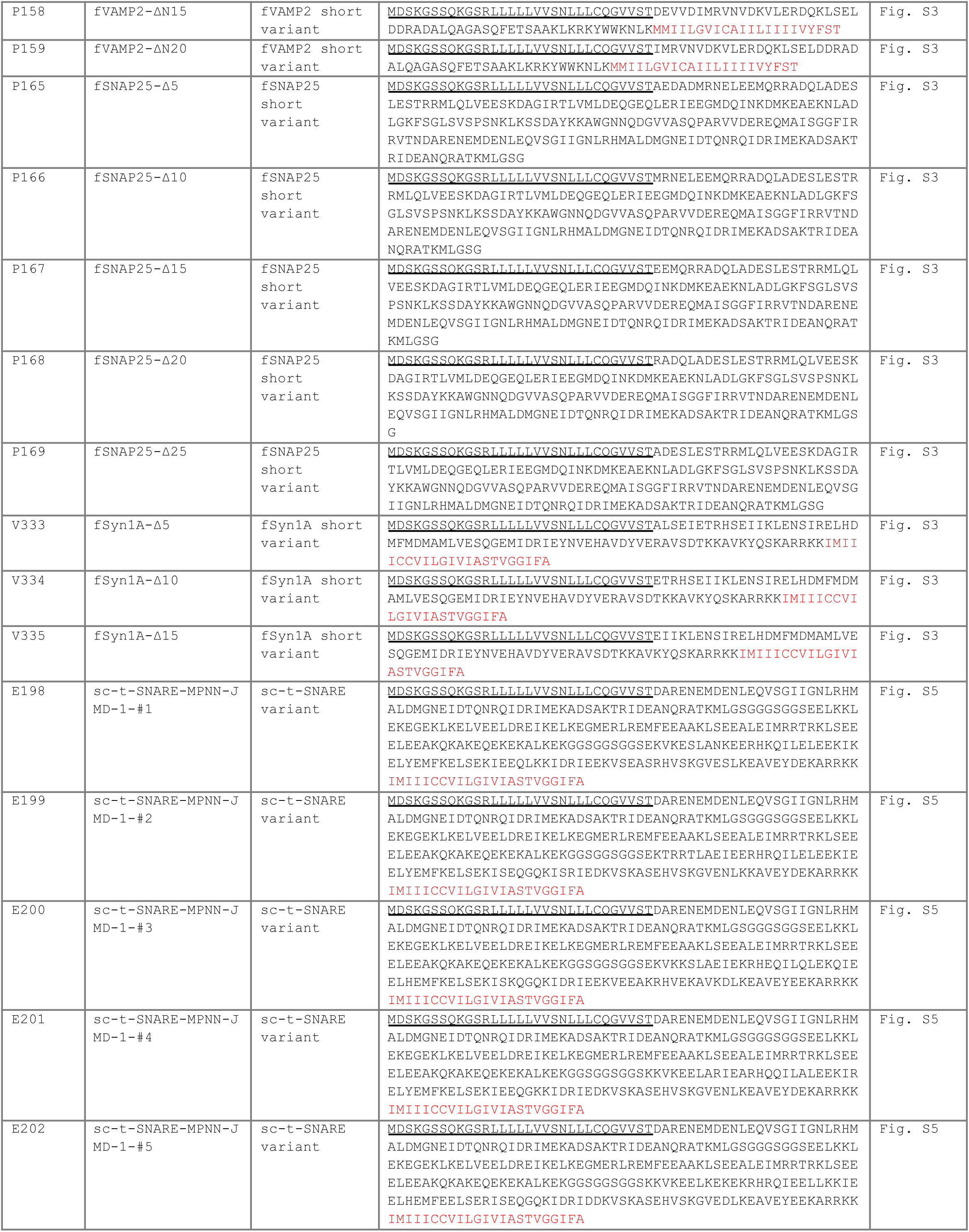
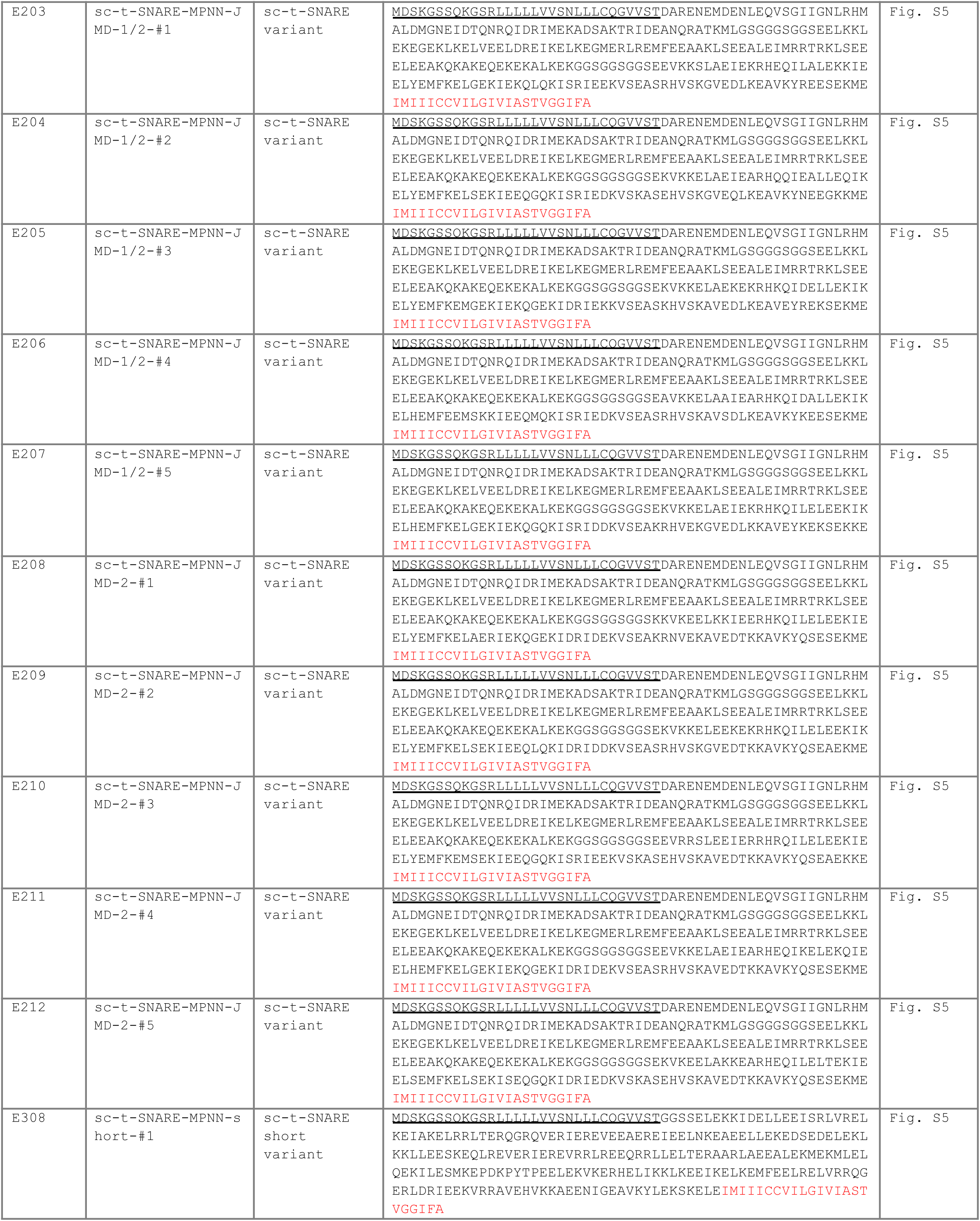
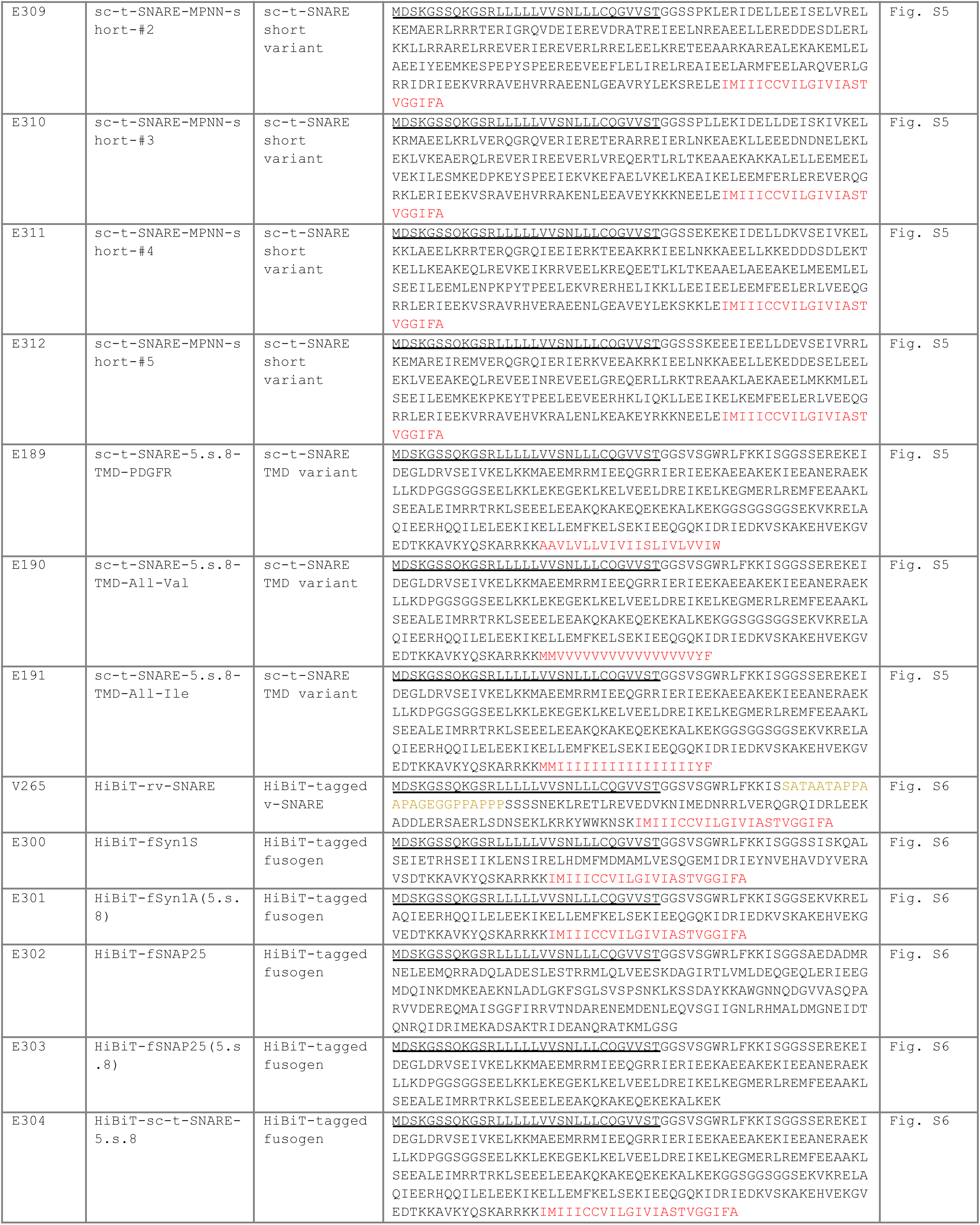
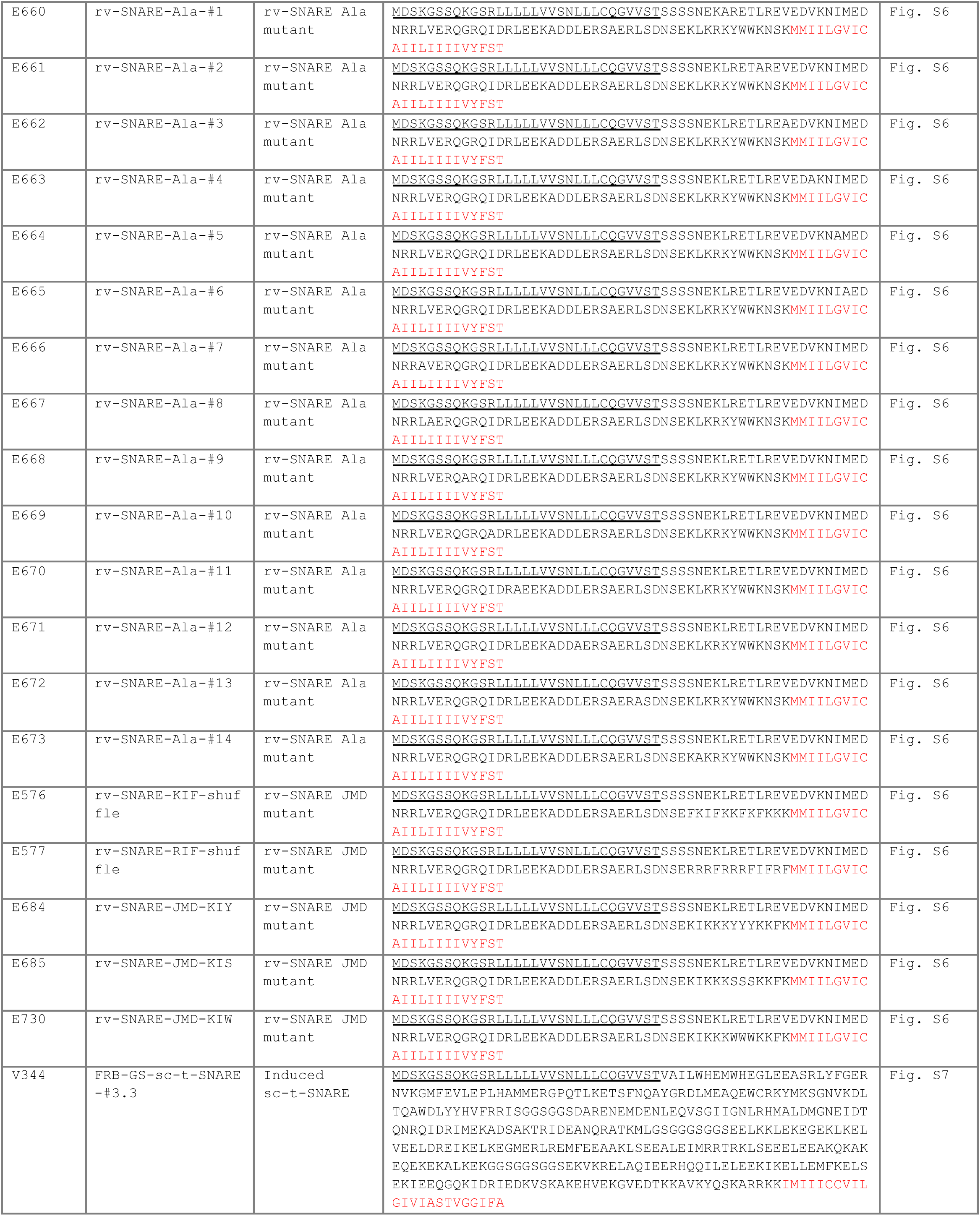
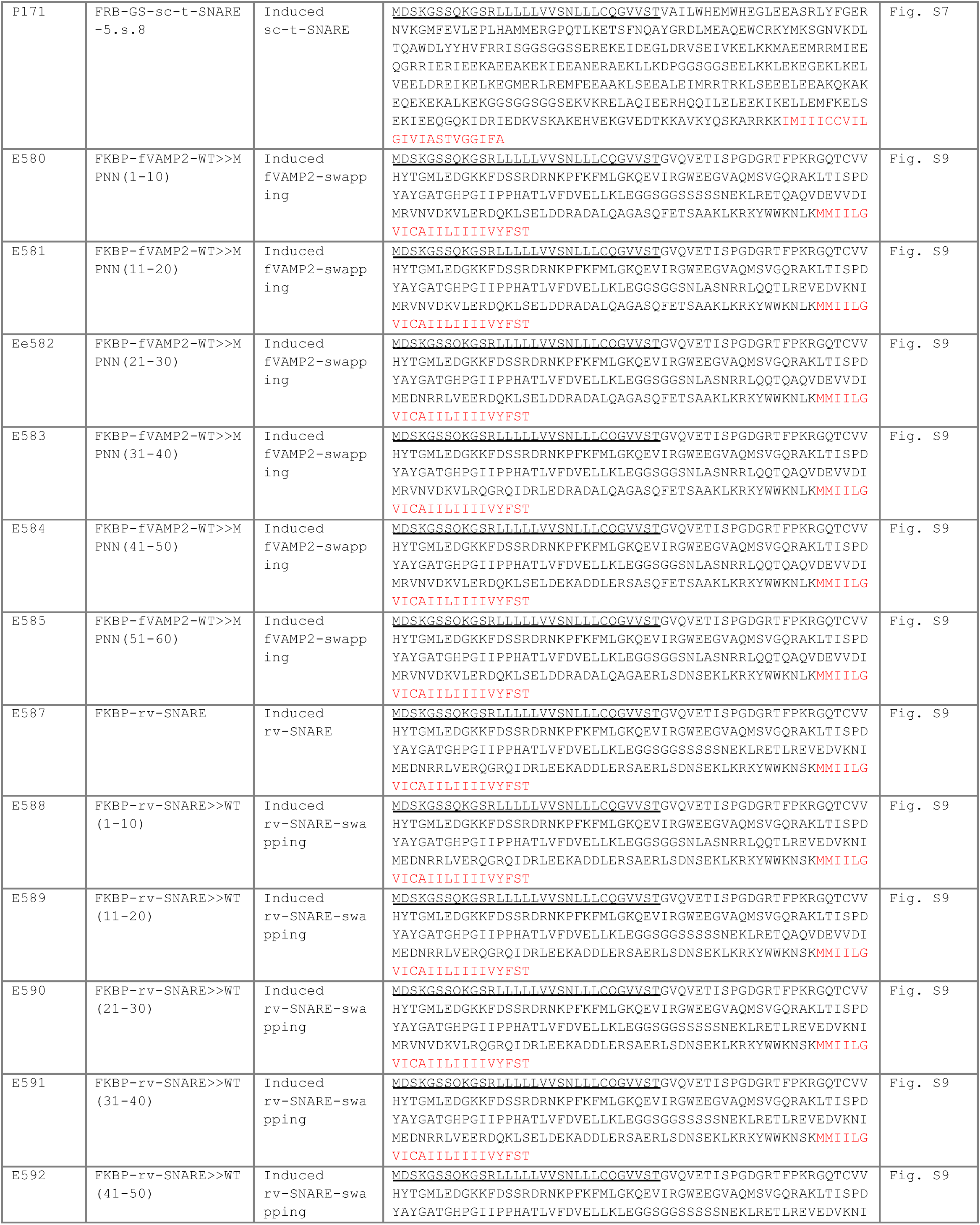
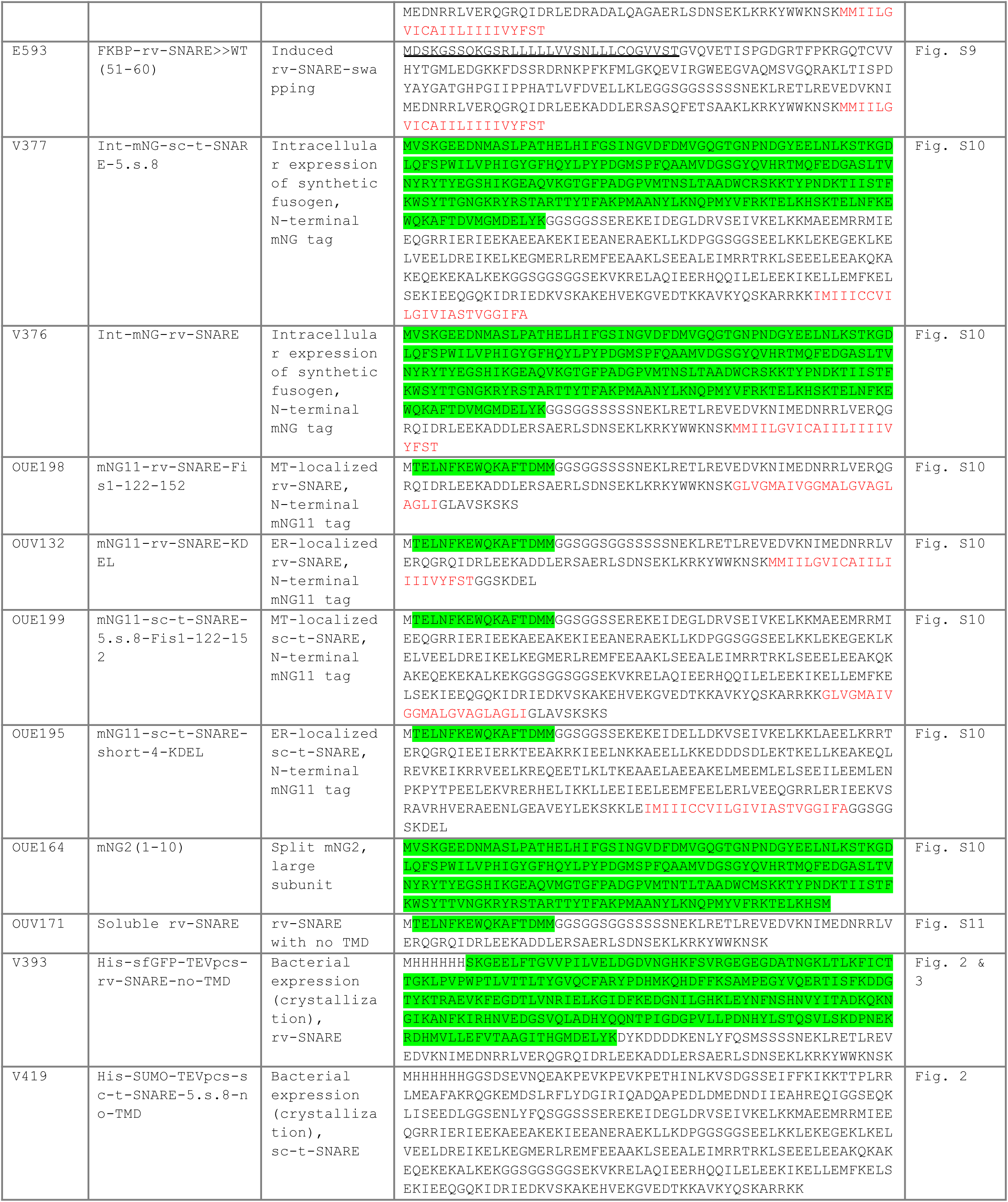
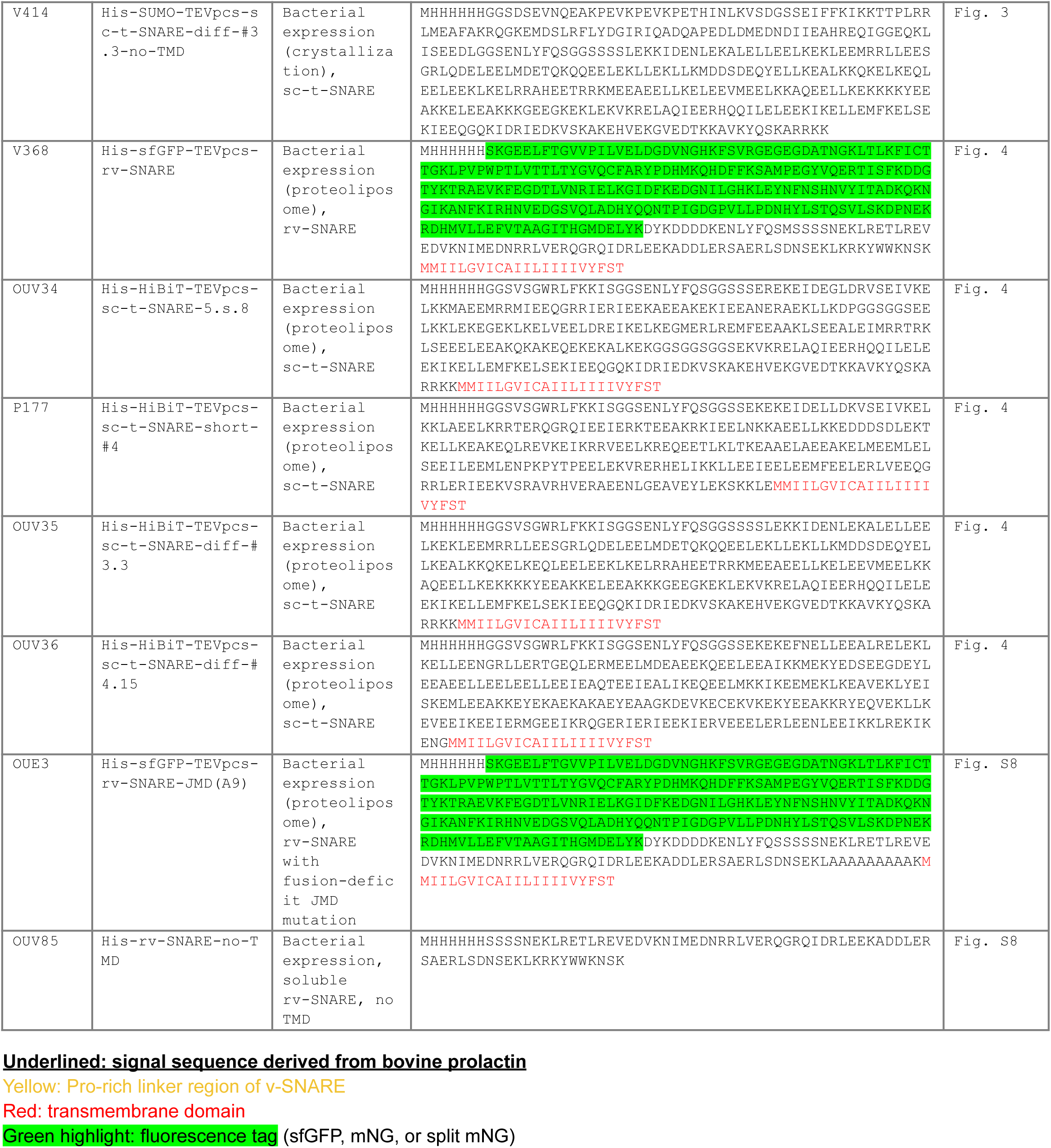
Protein sequences.

**Table S3.** Data collection and refinement statistics.

|  | rv-SNARE/sc-t-SNARE-diff-<br>#3.3<br>(PDB Code: 9Y5H) | rv-SNARE/sc-t-SNARE-5.s.8<br>complex (space group P4<sub>1</sub>2<sub>1</sub>2)<br>(PDB Code: 9Y5F) | rv-SNARE/sc-t-SNARE-5.s.8<br>complex (space group<br>P2<sub>1</sub>2<sub>1</sub>2<sub>1</sub>) (PDB Code: 9Y5G) |
| --- | --- | --- | --- |
| **Data collection\*** |  |  |  |
| Space group | P 2<sub>1</sub> 2<sub>1</sub> 2<sub>1</sub> | P 4<sub>1</sub> 2<sub>1</sub> 2 | P 2<sub>1</sub> 2<sub>1</sub> 2<sub>1</sub> |
| Cell dimensions |  |  |  |
| *a*, *b*, *c* (Å) | 49.92, 56.55, 130.41 | 47.09, 47.09, 334.30 | 43.18, 45.70, 341.12 |
| $\alpha$, $\beta$, $\gamma$ (°) | 90, 90, 90 | 90, 90, 90 | 90, 90, 90 |
| Resolution (Å) | 46.62 - 2.52 (2.62 - 2.52)<sup>†</sup> | 33.14 - 3.45 (3.78 - 3.45) | 33.34 - 2.02 (2.07 - 2.02) |
| *R*<sub>sym</sub> or *R*<sub>merge</sub> | 0.088 (0.941) | 0.174 (0.531) | 0.250 (2.143) |
| *I* / $\sigma$ | 13.1 (2.2) | 15.3 (8.6) | 6.1 (0.8) |
| Completeness (%) | 99.9 (100) | 99.8 (100) | 100 (100) |
| Redundancy | 6.4 (6.9) | 23.8 (26.1) | 9.0 (8.4) |
| **Refinement** |  |  |  |
| Resolution (Å) | 42.72 - 2.52 (2.62 - 2.52) | 33.30 - 3.45 (3.80 - 3.45) | 34.11 - 2.02 (2.07 - 2.02) |
| No. reflections | 12,993 (1,409) | 5,481 (1,300) | 40,812 (2,885) |
| *R*<sub>work</sub> / *R*<sub>free</sub> | 0.2360 (0.3288) / 0.2939<br>(0.4356) | 0.2267 (0.2196) / 0.2880<br>(0.2998) | 0.2274 (0.2995) / 0.2781<br>(0.3125) |
| No. atoms |  |  |  |
| Protein | 2,594 | 2,534 | 5,004 |
| Water | 30 | n/a | 302 |
| *B*-factors |  |  |  |
| Protein | 69 | 74 | 37 |
| Water | 61 | n/a | 36 |
| R.M.S. deviations |  |  |  |
| Bond lengths (Å) | 0.001 | 0.001 | 0.002 |
| Bond angles (°) | 0.290 | 0.320 | 0.350 |

\*Single crystal used for each dataset/structure.
<sup>†</sup>Values in parentheses are for the highest-resolution shell.

